# Multiple orthoflaviviruses secrete sfRNA in mosquito saliva to promote transmission by inhibiting MDA5-mediated early interferon response

**DOI:** 10.1101/2025.01.21.634113

**Authors:** Idalba Serrato-Pomar, Hacène Medkour, Louise Belleville, Kachaporn Jintana, Lauryne Pruvost, Norman Schneider, Mihra Tavadia, Jim Zoladek, Florian Rachenne, Zoé Roux, Quentin Narpon, Felix Rey-Cadilhac, Solena Rossi, Stephanie French, Cassandra Modahl, Carole Ginibre, Bethsabée Scheid, Wannapa Sornjai, Elliott F. Miot, Duncan R. Smith, Rodolphe Hamel, Oleg Medianikov, Dorothée Missé, Sébastien Nisole, Julien Pompon

**Affiliations:** MIVEGEC, Univ. Montpellier, IRD, CNRS, Montpellier, France; IHU Méditerranée Infection, Marseille, France; IRIM, Univ. Montpellier, CNRS, Montpellier, France; Liverpool School of Tropical Medicine, Liverpool, United Kingdom; Institute of Molecular Biosciences, Mahidol University, Bangkok, Thailand; Department of Clinical Microbiology and Applied Technology, Faculty of Medical Technology, Mahidol University, Nakhon Pathom, Thailand; Viral Vector Joint unit and Joint Laboratory, Mahidol University, Nakhon Pathom, Thailand; IRD, AP-HM, MEPHI, Aix Marseille University, Marseille, France; Department of Medical Technology, LMI PRESTO, Faculty of Associated Medical Sciences, Chiang Mai University, Chiangmai, Thailand; Centre Armand-Frappier Santé Biotechnologie, Institut National de la Recherche Scientifique (INRS), Laval, QC, Canada

**Author notes:** Authors contributed equally (Louise Belleville, Kachaporn Jintana, Lauryne Pruvost, Norman Schneider, Mihra Tavadia).

**Keywords:** Mosquito-borne diseases, transmission, extracellular vesicles, innate immunity, skin

## Abstract

Numerous orthoflaviviruses transmitted through the bites of different mosquito species infect more than 500 million people annually. Skin infection at the bite site represents a critical and conserved step in transmission and a deeper understanding of this process will promote the design of broad-spectrum interventions to address diverse orthoflavivirus health threats. Here, we identify and characterize a transmission-enhancing viral factor in mosquito saliva that is shared across orthoflaviviruses. Saliva from West Nile virus-infected *Culex* and Zika virus-infected *Aedes* contains a viral non-coding RNA, subgenomic flaviviral RNA (sfRNA), within lipid vesicles distinct from virions. Higher concentration of sfRNA in infectious saliva positively correlates with infection intensity in human cells and skin explants. Early sfRNA delivery into transmission-relevant skin cell types and human skin explant demonstrate that sfRNA is responsible for the infection enhancement. Co-inoculation of sfRNA in a mouse model of transmission enhanced skin infection and worsened disease severity, supporting the role of salivary sfRNA as a transmission-enhancer. Mechanistically, salivary sfRNA attenuates early interferon response in human skin cells and skin explants by disrupting MDA5 signaling. Our results, derived from two distinct orthoflaviviruses and supported by prior studies, establish salivary sfRNA as a pan-orthoflavivirus transmission-enhancing factor driven by a conserved viral non-coding RNA.

## Introduction

Numerous mosquito-borne orthoflaviviruses cumulatively infect half a billion people annually, resulting in an estimated 150,000 fatalities and economic losses exceeding €9 billion ^1^. Given the wide geographic distribution of *Aedes* and *Culex* mosquito vectors, nearly the entire human population faces the risk of infection ^2,3^. Furthermore, predictive models considering increasing urbanization, global transportation, and climate change anticipate exacerbation and broadening of the associated public health, economic, and social burdens ^4^. More concerning is the very likely emergence of yet-unknown mosquito-borne orthoflaviviruses in the near future, further complicating the landscape of orthoflavivirus diseases that necessitate targeted interventions ^1^. However, safe and effective interventions to protect against orthoflaviviral diseases are lacking. While vector mosquito control stands as the most widely implemented strategy, its effectiveness in preventing epidemics is only moderate ^5^. Scaling vector control approaches to cover extensive territories is also challenging, and the efficacy is hindered by the rapid development of insecticide resistance ^6^. Additionally, there is no curative treatment ^7^, and the available vaccines pose notable safety concerns ^8,9^. A promising approach to confront the rising threat posed by multiple orthoflaviviruses is the design of pan-orthoflavivirus interventions, achieved through the identification of targets that are conserved across orthoflaviviruses.

The transmission of mosquito-borne orthoflaviviruses occurs during a mosquito bite ^10^ (although alternative transmission routes such as sexual and maternal transmissions are possible ^1^), when infectious virions present in the saliva are deposited in the epidermis and dermis ^11,12^. Subsequent productive infection of the skin is essential for transmission ^13,14^. However, infection initiation is hindered by a potent antiviral response in the skin ^15,16^. Viral double-stranded RNA (dsRNA), an intermediate of viral replication, is detected by retinoic acid-inducible gene I (RIG-I), melanoma differentiation-associated gene 5 (MDA5) or toll-like receptor 3 (TLR3) ^17^. These three viral RNA sensing branches separately and synergistically activate biochemical cascades that converge onto the phosphorylation of interferon regulatory factor-3 (IRF3) and/or IRF7, which trigger the expression of type I interferon (IFN). IFN then triggers an antiviral state by inducing the expression of IFN-stimulated genes (ISGs) with anti-orthoflaviviral properties ^18,19^. Robust evidence supports that mosquito saliva modulates the antiviral innate immune response in the skin to promote orthoflavivirus infection ^13,20–22^. Nonetheless, identification of these anti-immune transmission-enhancing factors remains partial.

Recently, we discovered a mosquito saliva factor that augments saliva infectivity for dengue virus (DENV), the most prevalent *Orthoflavivirus* ^23^. DENV secretes a non-coding subgenomic orthoflaviviral RNA (sfRNA) with anti-immune properties ^24–26^ within salivary lipid exosomes to enhance saliva infectivity ^27^. Nonetheless, our prior study did not provide evidence of an effect on transmission and lacked a mechanistic understanding. SfRNA biogenesis results from the incomplete degradation of orthoflaviviral genomic RNA (gRNA) by 5’-to-3’ host exoribonucleases, which stall on 3’UTR structures, called exoribonuclease-resistant RNA (xrRNA) ^28^. Intriguingly, the process of sfRNA biogenesis is conserved across orthoflaviviruses ^29–32^, raising the hypothesis that salivary sfRNA secretion occurs for multiple orthoflaviviruses.

West Nile virus (WNV) and Zika virus (ZIKV) rank among the most important orthoflaviviruses in terms of global public health burden. WNV is the most widely distributed *Orthoflavivirus* ^33^ and elicits symptoms potentially resulting in severe neurological manifestations leading to mortality or persistent sequelae ^34^. Since 2000, WNV has infected an estimated 7 million people, causing 2,700 deaths in the USA ^35^, and is annually responsible for over 100 fatalities in the EU ^36^. ZIKV, acknowledged as a pathogen of international concern by the World Health Organization (WHO) in 2016, has infected more than 2 million people since its emergence ^37,38^. While ZIKV symptoms typically present as mild, the infection poses substantial risks of conge ital Zika syndrome in newborns of infected mothers and Guillain-Barré syndrome in infected adults. Importantly, WNV and ZIKV exhibit distinct transmission dynamics with WNV transmitted by various *Culex* mosquitoes, notably *Cx. quinquefasciatus* ^39^, whereas ZIKV is primarily disseminated by *Aedes* mosquitoes, predominantly *Ae. aegypti* ^40,41^.

Here, using WNV and ZIKV as *Orthoflavivirus* models, we investigated whether sfRNA secretion in mosquito saliva represents a pan-orthoflavivirus mechanism for enhancing transmission and elucidated how salivary sfRNA increases transmission. First, we detected sfRNA in the saliva of *Culex* mosquitoes infected with WNV and *Aedes* mosquitoes infected with ZIKV. Second, we found that salivary sfRNA from WNV-infected *Culex* and ZIKV-infected *Aedes* is packaged within lipid vesicles. Third, combining approaches with transmission-relevant human skin cell types, human skin explants and a mouse model of transmission, we demonstrated that salivary sfRNA enhances *Orthoflavivirus* transmission. Finally, we showed that early sfRNA exposure suppresses the initial IFN response by disrupting MDA5-mediated signaling.

## Results

### SfRNA is secreted in saliva from WNV-infected *Culex* and ZIKV-infected *Aedes* mosquitoes

WNV can produce up to four sfRNA species by stalling host exoribonucleases at distinct xrRNA structures ^42^. To identify the sfRNA species in *Cx. quinquefasciatus*, we performed Northern blot on mosquitoes orally infected with WNV and observed a major band above 500 nt (Fig. 1a), as previously reported in *Cx. pipiens* mosquitoes ^43^ and corresponding to the predicted sfRNA1 species at 528 nt. We also noted a fainter band above 400 nt, which does not align with any predicted xrRNA, but was previously observed in monkey Vero cells ^43^. We then collected saliva and salivary glands (SG) – the site of saliva production - from *Culex* mosquitoes 10 days post oral infection and using absolute quantification (S1a,b Fig.) detected WNV gRNA in 80% of SG with a geometric mean of 1.95 × 10^3^ copies per infected sample and in 60% of saliva with a geometric mean of 1.97 × 10^2^ copies per infected sample (Fig. 1b). Using an optimized subtractive RT-qPCR protocol for absolute quantification of sfRNA (Supplementary Materials; S2a-d and S3a,b Fig.), we detected sfRNA in 93% of gRNA-positive SG with a geometric mean of 1.96 × 10^6^ per infected sample and in 100% of gRNA-positive saliva with a geometric mean of 4.15 × 10^5^ copies per infected sample (Fig. 1c). To normalize sfRNA to the infection level (estimated by gRNA level), we calculated the ratio of sfRNA:gRNA and observed ratios in SG and saliva at 1,005 and 1,431, respectively (Fig. 1d). To gain deeper insights into sfRNA secretion mechanism, we further analysed the data of salivary sfRNA secretion. In SG, we observed a positive correlation between sfRNA and gRNA copies (Fig. 1e), indicating a relationship between sfRNA and its precursor in this organ, coherent with sfRNA biogenesis occuring within the SG. In contrast, in saliva, we identified a negative correlation between sfRNA and gRNA levels (Fig. 1f). These opposing correlations between gRNA and sfRNA in the SG and saliva are consistent with the two viral RNAs being secreted through distinct mechanisms. By analyzing paired SG and saliva samples from the same mosquito, we further examined correlations between the two compartments. The quantities of gRNA and sfRNA in saliva remained relatively constant and showed no significant association with their levels in the SG (S4a,b Fig.). Collectively, our findings provide evidence that WNV secretes sfRNA into *Culex* mosquito saliva and underscore the pivotal role of the SG in the production of expectorated sfRNA.

**Fig. 1.**
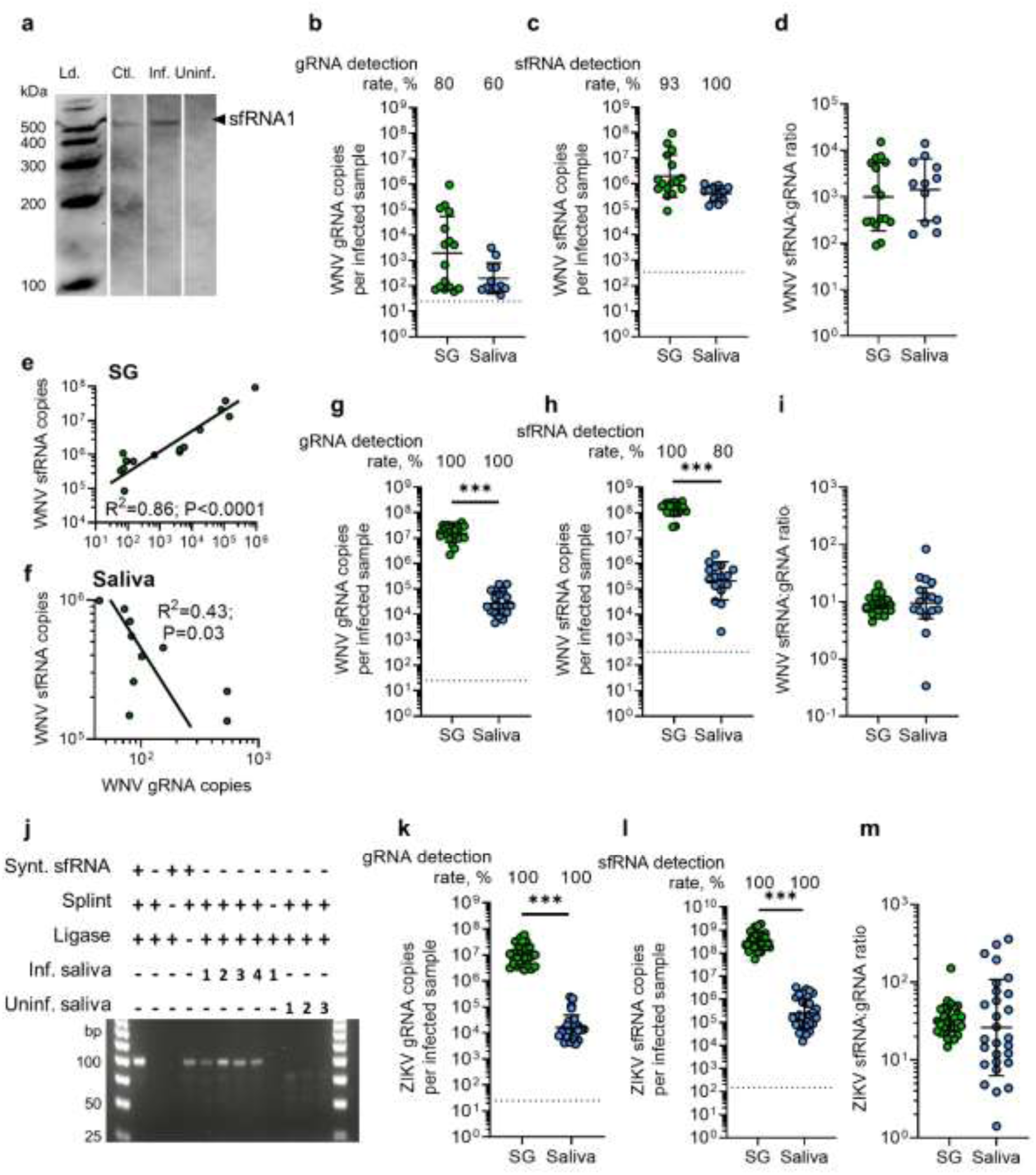
sfRNA is secreted in saliva from WNV-infected *Culex* and ZIKV-infected *Aedes* mosquitoes. **a** Detection of WNV sfRNA in orally infected *Culex* mosquitoes by Northern blot. Ld., RNA ladder; Ctrl., *in vitro* transcribed folded WNV sfRNA1; Inf., WNV-infected *Culex* mosquitoes; Uninf., mock-infected *Culex* mosquitoes. **b-d** Quantification of gRNA (b), sfRNA (c), and the ratio of sfRNA:gRNA (d) in salivary glands (SG) and saliva from *Culex* mosquitoes orally infected with WNV. **e-f** Correlations between gRNA and sfRNA quantity in SG (e) and saliva (f) from *Culex* orally infected with WNV. **g-i** Quantification of gRNA (g), sfRNA (h), and the ratio of sfRNA:gRNA (i) in SG and saliva from *Culex* mosquitoes micro-injected with WNV. **j** Direct detection of WNV sfRNA using SL-RT-PCR in saliva from WNV micro-injected mosquitoes. Number for infected and uninfected saliva indicate sample number. **k-m** Quantification of gRNA (k), sfRNA (l), and the ratio of sfRNA:gRNA (m) in SG and saliva from *Aedes* mosquitoes micro-injected with ZIKV. Each point represents one pair of SG or saliva collected from one mosquito. b-d, g-i, k-m Lines show geometric means ± 95% C.I. Dotted lines indicate LoD. sfRNA detection rate was calculated among gRNA-positive samples. ***, p < 0.001 according to T-test.

To further support the role of the SG, we bypassed midgut infection by inoculating WNV into the thorax of *Culex* mosquitoes and collected SG and saliva at 7 days post-infection (dpi), i.e. earlier than after oral infection due to the shortened extrinsic incubation period. Direct injection resulted in 100% of SG and saliva samples positive for gRNA, with geometric means of 1.57 × 10^7^ and 2.72 × 10^4^ copies per sample, respectively (Fig. 1g). SfRNA was detected in 100% of infected SG, with a geometric mean of 1.40 × 10^8^ copies per sample, and in 80% of infected saliva, with a geometric mean of 2.09 × 10^5^ copies per sample (Fig. 1h). This corresponded to sfRNA:gRNA ratios at 8.9 in SG and 9.5 in saliva (Fig. 1i). As with samples from orally-infected mosquitoes, we observed a strong positive correlation between gRNA and sfRNA in SG (S5a Fig.); however the correlation was maintained in saliva (S5b Fig.), likely reflecting conditions associated with micro-injection. Collectively, these results indicate that sfRNA secretion is an intrinsic feature of salivary gland infection rather than dependent on the route of viral entry.

To ascertain the secretion of sfRNA in saliva, we applied two additional methods. First, we conducted a NB on a pool of hundreds of mosquito saliva samples and detected a band of the expected size, but very faint (S6 Fig.). Second, we designed a direct molecular detection assay for sfRNA by applying splinted-ligation reverse-transcription PCR (SL-RT-PCR), as for mRNA decay quantification^44^. This method overcomes a major limitation in the field, where direct detection of sfRNA in low-input biological samples has remained challenging. Briefly, a DNA splint complementary to both an anchor RNA and the sfRNA sequences brings the two RNA molecules into close proximity, enabling T4 DNA ligase-mediated ligation of the monophosphorylated 5’ end sfRNA to the anchor RNA. After splint DNA degradation, the resulting chimeric RNA fragment is specifically RT-PCR-amplified using primers annealing onto the anchor RNA and sfRNA sequences. We validated the detection method with a syntentic 5’ end monophosphorylated fragment of sfRNA, amplifying a band of 98 bp as expected, and showed the absence of amplification without the sfRNA target or the splint fragment (Fig. 1j, lanes 1-3), although amplification occurred without ligase (Lane 4), indicating ligase-independent ligation. We then applied the SL-RT-PCR to four pools of 10 saliva (samples 1-4) from WNV-infected *Culex* mosquitoes, containing 2.25 × 10^5^, 4.4 × 10^5^, 1.45 × 10^6^ and 9.7 × 10^4^ sfRNA copies for samples 1-4, respectively, as determined by substractive RT-qPCR method (S3 Fig.) and detected sfRNA in all samples (Fig. 1j, lanes 5-8). We repeated the molecular assay on the saliva pool sample 1 but omitting the splint fragment to confirm specificity (Fig. 1j, lane 9). Finally, three pools of 10 saliva samples from uninfected mosquitoes did not result in amplification (Fig. 1j, lanes 10-12), confirming the specificity of the direct molecular detection of sfRNA and thereby the presence of sfRNA in saliva.

To determine whether sfRNA secretion in saliva is conserved among orthoflaviviruses transmitted by different mosquito genera, we inoculated *Aedes* mosquitoes with ZIKV and collected SG and saliva at 7 dpi. In whole mosquitoes, NB showed the presence of a single sfRNA band at the expected size (S7 Fig.), as previously detected in *Ae. aegypti* ^45^. ZIKV gRNA was detected in 100% of SG and in 100% of saliva samples, with geometric means of 1.00 × 10^7^ and 1.63 × 10^4^ copies per sample, respectively (Fig. 1k; S8a,b Fig.). We quantified ZIKV sfRNA using primers that annealed to both previously identified sfRNA species in *Ae. aegypti* ^46^ and an optimized sfRNA absolute quantification method (Supplementary materials; S9a-c and S10a,b Fig.). ZIKV sfRNA was detected in 100% of infected SG, with a geometric mean of 3.22 × 10^8^ copies per sample, and in 100% of infected saliva, with a geometric mean of 2.47 × 10^5^ copies per sample (Fig. 1l). This resulted in sfRNA:gRNA ratios of 32.2 in SG and 26.1 in saliva (Fig. 1m). As for WNV-infected *Culex*, ZIKV gRNA and sfRNA correlations were different between SG and saliva with a weaker correlation in saliva (S11a,b Fig.). Collectively, the findings demonstrate that ZIKV, another *Orthoflavivirus*, secretes sfRNA in the saliva of *Aedes* mosquitoes, indicating that sfRNA secretion in saliva is a conserved mechanism among orthoflaviviruses vectored by different mosquito species.

### Salivary sfRNAs from WNV-infected *Culex* and ZIKV-infected *Aedes* are within lipid vesicles

In our previous study, we provided biochemical and microscopic evidence that DENV sfRNA is expectorated within lipid vesicles distinct from virus particles ^23^. To determine whether this phenomenon is shared by other orthoflaviviruses, we assessed the nuclease resistance of salivary sfRNA from WNV-inoculated *Culex* mosquitoes following detergent-based disruption of lipid membranes (Fig. 2a). We showed that micrococcal nuclease (MNase) degraded *in vitro* transcribed WNV sfRNA (Fig. 2b) (RNase A/T1 was inefficient in degrading WNV sfRNA) and a viral RNA control fragment (S12a Fig.) in the presence of Triton-X 100 detergent and uninfected *Culex* saliva. Consistent with salivary gRNA and sfRNA being enclosed within lipid particles, neither of the viral RNAs were degraded by MNase unless pre-treated with Triton X-100 (Fig. 2c,d). To account for variability in the initial amounts of sfRNA and gRNA between the saliva pools, we normalized the RNA quantities within each saliva pool and observed consistent trends (S13a-c Fig.). We then reasoned that if sfRNA and gRNA are enclosed in separate lipid particles, they would exhibit different sensitivities to the detergent treatment. Leveraging the inherent variability between biological replicates – potentially due to mechanical stresses from pipetting and liquid handling^47^ – we found a lack of correlation between sfRNA and gRNA levels after detergent and MNase treatments (Fig. 2e). This observation suggests that sfRNA is encapsulated in distinct lipid particles, separate from those containing gRNA.

**Fig. 2.**
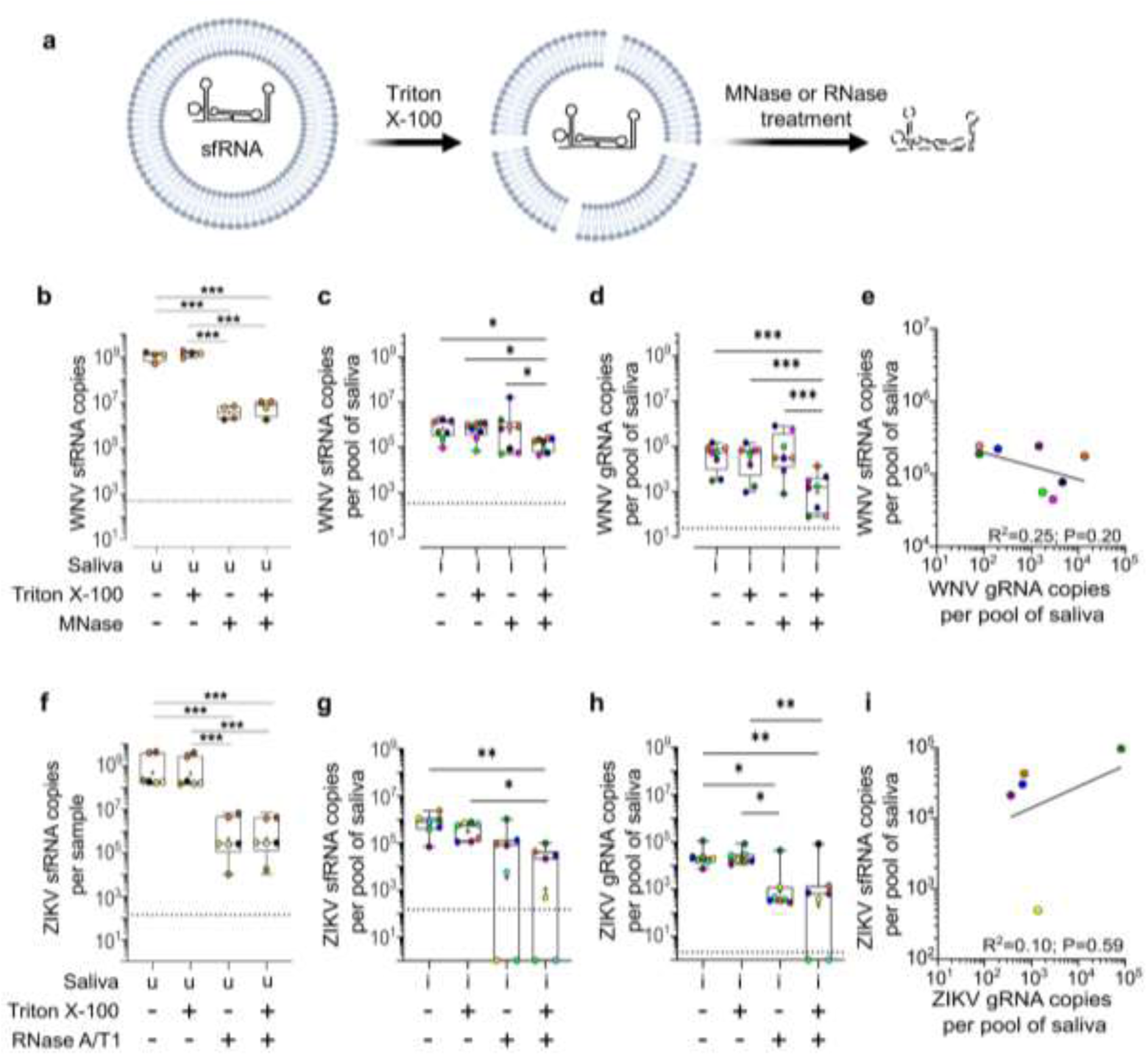
sfRNA in WNV-infected *Culex* saliva and ZIKV-infected *Aedes* saliva is protected from nuclease degradation by a detergent sensitive layer. **a** sfRNA inside a lipid bilayer is sensitized to nuclease degradation by Triton X-100 treatment. **b-d** Sensitivity to triton X-100 and Micrococcal nuclease (MNase) treatment for *in vitro*-transcribed WNV sfRNA diluted in uninfected *Culex* saliva (b), for salivary WNV sfRNA (c) and for salivary WNV gRNA (d). Saliva was collected from WNV-inoculated *Culex* mosquitoes. **e** Correlation between sfRNA and gRNA quantities in WNV-infected *Culex* saliva after triton X-100 and MNase treatments. **f-h** Sensitivity to triton X-100 and RNase A/T1 treatment for *in vitro*-transcribed ZIKV sfRNA diluted in uninfected *Aedes* saliva (f), for salivary ZIKV sfRNA (g) and for salivary ZIKV gRNA (h). Saliva was collected from ZIKV-inoculated *Aedes* mosquitoes. **i** Correlation between sfRNA and gRNA quantities in ZIKV-infected *Aedes* saliva after triton X-100 and RNase A/T1 treatments. Boxplots indicate median ± lower and higher quartiles. Dots indicate saliva pool repeats. The dot colors indicate the different pools of saliva. u, uninfected saliva; i, infected saliva from the corresponding mosquito. *, p < 0.05; **, p < 0.01; ***, p < 0.001 according to T-test.

We repeated the nuclease resistance assay using saliva from ZIKV-inoculated *Aedes* mosquitoes. After confirming the degradation of *in vitro*-transcribed ZIKV sfRNA (Fig. 2f) and viral control RNA (S12b Fig.) by RNase A/T1 (RNase), we observed that salivary sfRNA was partially protected from RNase degradation unless pretreated with a detergent (Fig. 2g). Surprisingly, ZIKV sfRNA and gRNA in saliva were partially degraded by RNase both with and without detergent pre-treatment (Fig. 2h), potentially due to unintended physical stress on the virions during the one-day-long process from saliva collection to RNase treatment. Nonetheless, leveraging this unexpected sensitization, we observed a discrepancy between sfRNA and gRNA quantities (Fig. 2g,h), consistent with the two RNA fragments being enclosed in different particles. Similar trends were observed after normalizing RNA quantities within saliva pools (S13d-f Fig.). Finally, we noted the absence of correlation between salivary sfRNA and gRNA in samples treated with both detergent and RNase (Fig. 2i) – especially clear for two samples with detectable level of sfRNA and non-detectable level of gRNA. Altogether, the analysis of saliva from two different mosquito genera infected with different orthoflaviviruses suggest the conservation of sfRNA secretion within salivary lipid vesicles across multiple orthoflaviviruses.

### Salivary sfRNA increases WNV transmission

To elucidate the role of salivary sfRNA in transmission, we infected *Culex* mosquitoes with WNV by oral feeding, which introduces greater variability in the salivary sfRNA:gRNA ratio compared to thoracic inoculation (Fig. 1). Saliva pools were collected and categorized based on their sfRNA:gRNA ratio into low (ratio < 15), moderate (15 < ratio < 350), or high (350 < ratio) sfRNA concentrations (Fig. 3a; S1 Table). To validate the use of the saliva-collection media as an inoculum, we determined that Erioglaucine, used as a proxy for salivation in the saliva-collection media, did not affect cell viability (S14 Fig.). To support the normalization of saliva inoculum dose based on gRNA quantities, we showed that gRNA quantities were proportional to saliva infectiousness as measured by particle forming unit (PFU) (R² = 0.98; p < 0.001; S15a Fig.) and that sfRNA concentration was not associated with the gRNA-to-PFU (PFU) ratio (S15b Fig.), a measure for infectivity per virion. Saliva from the different sfRNA concentration groups were then used to infect Huh7 human hepatocyte cells, using volumes containing the same number of gRNA copies (S1 Table). Total saliva volumes were normalized with uninfected saliva. Quantification of WNV infection at 24 and 48 hours post-infection (hpi) showed a progressive increase in infection with ascending categories of salivary sfRNA concentration (Fig. 3b,c) and a positive correlation with sfRNA:gRNA ratio (Fig. 3d). We repeated the saliva infection experiments with human skin explants (Fig. 3a). Injection of saliva containing higher sfRNA concentrations (S2 Table) led to increased WNV infection at 24hpi (Fig. 3e,f).

**Fig. 3.**
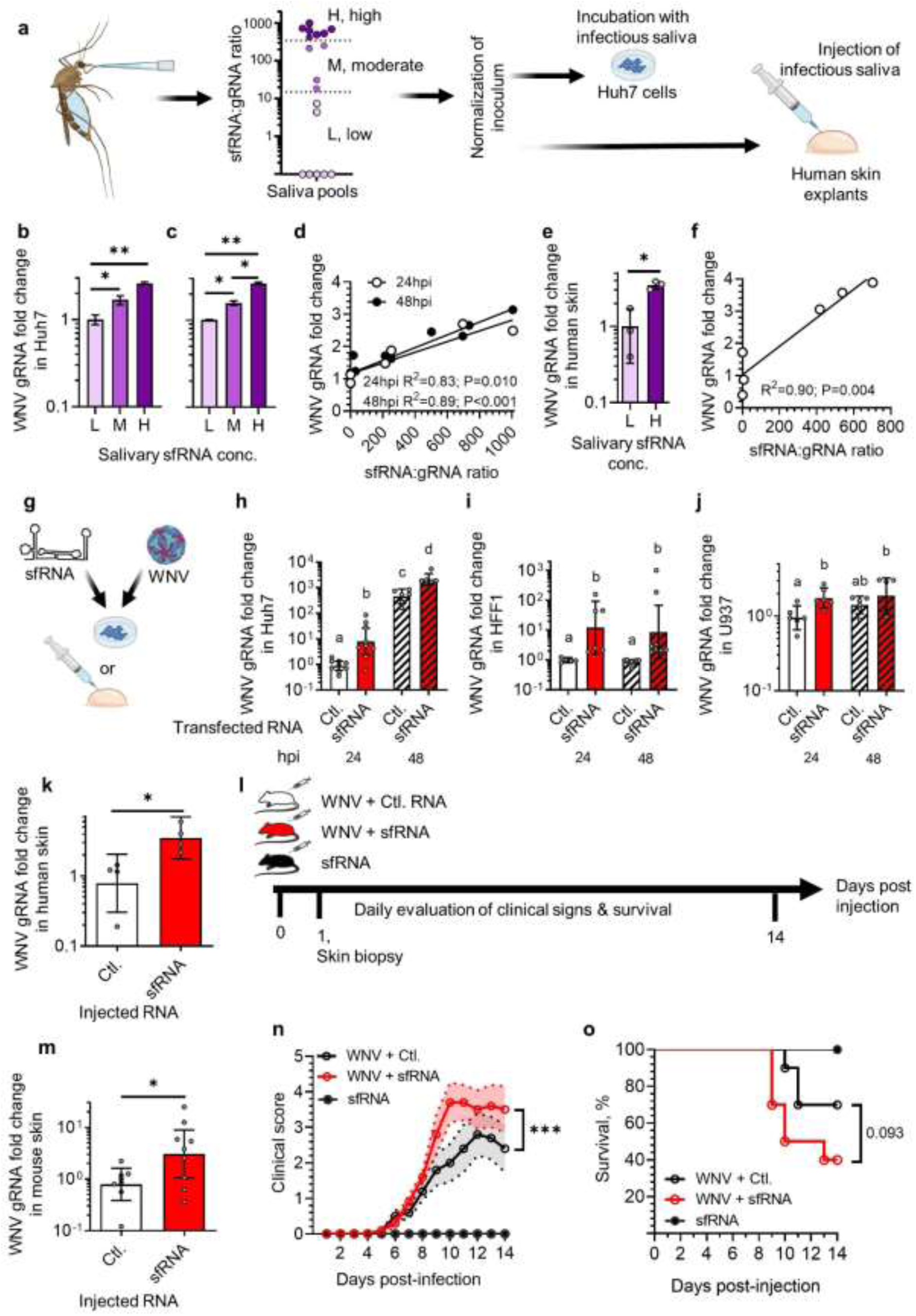
| SfRNA in saliva increases WNV transmission. **a** Pools of saliva were collected, categorized according to sfRNA:gRNA ratio and used to inoculate cells or human skin explants. **b-c** WNV infection in Huh7 cells at 24 (b) and 48 hpi (c) with low (L), moderate (M) and high (H) concentration of salivary sfRNA. Infection levels were normalized to L within time points. N per category, 2. **d** Correlation between infection intensity (i.e., WNV gRNA fold change) in Huh7 and sfRNA concentration (i.e., sfRNA:gRNA ratio) in the corresponding saliva used for infection. **e** WNV infection in skin explants at 24 h post injection with low and high salivary sfRNA concentration. **f** Correlation between infection intensity in skin explants and sfRNA concentration in the corresponding saliva used for infection. **g** Cells and skin explants were transfected and injected, respectively, with sfRNA or Control RNA (Ctl.) prior WNV infection. **h-j** WNV infection at MOI 0.005 in Huh7 (h), HFF1 (i) and U937 (j) cells at 24 and 48 hpi with WNV post sfRNA transfection. **k** WNV infection in skin explants at 24 hpi with WNV post sfRNA injection. **l** Mice were intradermally co-injected with WNV and sfRNA or Ctl. RNA. SfRNA alone was injected as control. Skin biopsies were collected at day 1 and clinical signs and survival were recorded daily. N, 14. **m** WNV infection in mouse skin at 24 h post injection. **n-o** Clinical signs (n) and survival (o) for mice co-injected with WNV and either sfRNA or Ctl. RNA. b, c, e Bars show mean ± s.e.m. h-j, k, m Bars show geometric mean ± 95% C.I. n Lines show mean ± s.e.m. Dots indicate repeats. Different letters show significant differences and *, p < 0.05; **, p < 0.01 according to post hoc Fisher’s LSD test, T-test or general linear mixed model. h-j, Different letters indicate statistical differences according to Fisher’s LDS test.

We next investigated whether early presence of sfRNA enhances infection. First, we transfected Huh7 cells with *in vitro*-transcribed sfRNA shortly before WNV infection (Fig. 3g). Intracellular levels of sfRNA or a size-matched viral RNA control were comparable after transfection, averaging ∼10^8^ copies per 250,000 cells (S16a Fig.). To achieve a sfRNA:gRNA ratio approximating that observed in mosquito saliva (Fig. 1d), we employed a low inoculum (MOI 0.005), corresponding to 1,250 PFU. Based on a gRNA:PFU ratio of 100 - commonly observed in virus preparations from mosquito cells ^48^ - this inoculum was estimated to contain 125,000 gRNA copies, resulting in a high sfRNA:gRNA ratio of 800. We observed that sfRNA transfection increased WNV infection at 24 and 48 hpi (Fig. 3h). Second, as orthoflaviviruses are thought to initially infect skin fibroblasts and monocytes following a mosquito bite ^10,49^, we repeated WNV infection post sfRNA transfection in HFF1 fibroblasts and U937 monocytes. While intracellular levels of transfected sfRNA and control RNA were equivalent (S16b,c Fig.), infection was increased after sfRNA transfection at 24 and 48 hpi in both cell types, although the increase was lower in U937 (Fig. 3i,j). To rule out biases associated with low MOI - where few cells are initially infected – and the use of high sfRNA:gRNA ratio, we repeated the experiments with a higher MOI of 0.5 in all three cell types, resulting in an estimated very low sfRNA:gRNA ratio of 8. Notably, the pro-viral function of sfRNA was maintained under these higher inoculum conditions (S17a-c Fig.). Third, we injected a mixture of WNV with either sfRNA or the control RNA into human skin explants (Fig. 3g) and observed that co-injection with sfRNA enhanced infection at 24 hpi (Fig. 3k).

Finally, we used a mouse model of transmission to evaluate the effect of sfRNA on infection and disease outcomes. Mice were intradermally injected with a small volume containing WNV and either sfRNA or the control RNA (Fig. 3l). As a non-infected control, mice were injected with sfRNA alone. At one day post injection (dpi), infection at the injection site in the skin was increased by co-injection with sfRNA (Fig. 3m). At 4 dpi, we confirmed successful systemic infection by detecting RNAemia (S18 Fig.), although this alone does not inform about disease severity ^50–52^. Over a longer observation period, we found that sfRNA co-injection aggravated disease severity (Fig. 3n), increased weight loss (S19 Fig.) and reduced mouse survival (Fig. 3o). Altogether, these results – obtained using infectious saliva with variable sfRNA concentrations and *in vitro*-transcribed sfRNA to mimic salivary delivery in *in vitro*, *ex vivo* and *in vivo* models - demonstrate that salivary sfRNA enhances transmission, worsening disease outcomes.

### Salivary sfRNA mitigates the IFN response

To decipher how WNV sfRNA impacts the innate immune response, we quantified expression of *IFN-β* and three ISGs, namely *CXCL10*, *MX1* and *IFI6*, in the Huh7 cell samples previously infected with saliva containing different sfRNA concentrations (same samples as in Fig. 3b-d). At 24 hpi, we observed a robust induction of *IFN-β* and the three ISGs in all samples treated with infectious saliva compared to mock-infected control (Fig. 4a-d). However, the levels of innate immune gene induction were similar across saliva samples irrespective of salivary sfRNA concentrations. Since cells infected with saliva containing higher sfRNA concentrations had higher viral loads (Fig. 3b) and previous studies have shown a correlation between viral load and intensity of IFN induction ^53,54^, these findings suggest that higher salivary sfRNA dampened the innate immune response at 24 hpi. In contrast, at 48 hpi, the expression of all four IFN-related genes correlated with the infection levels (Fig. 4a-d).

**Fig. 4.**
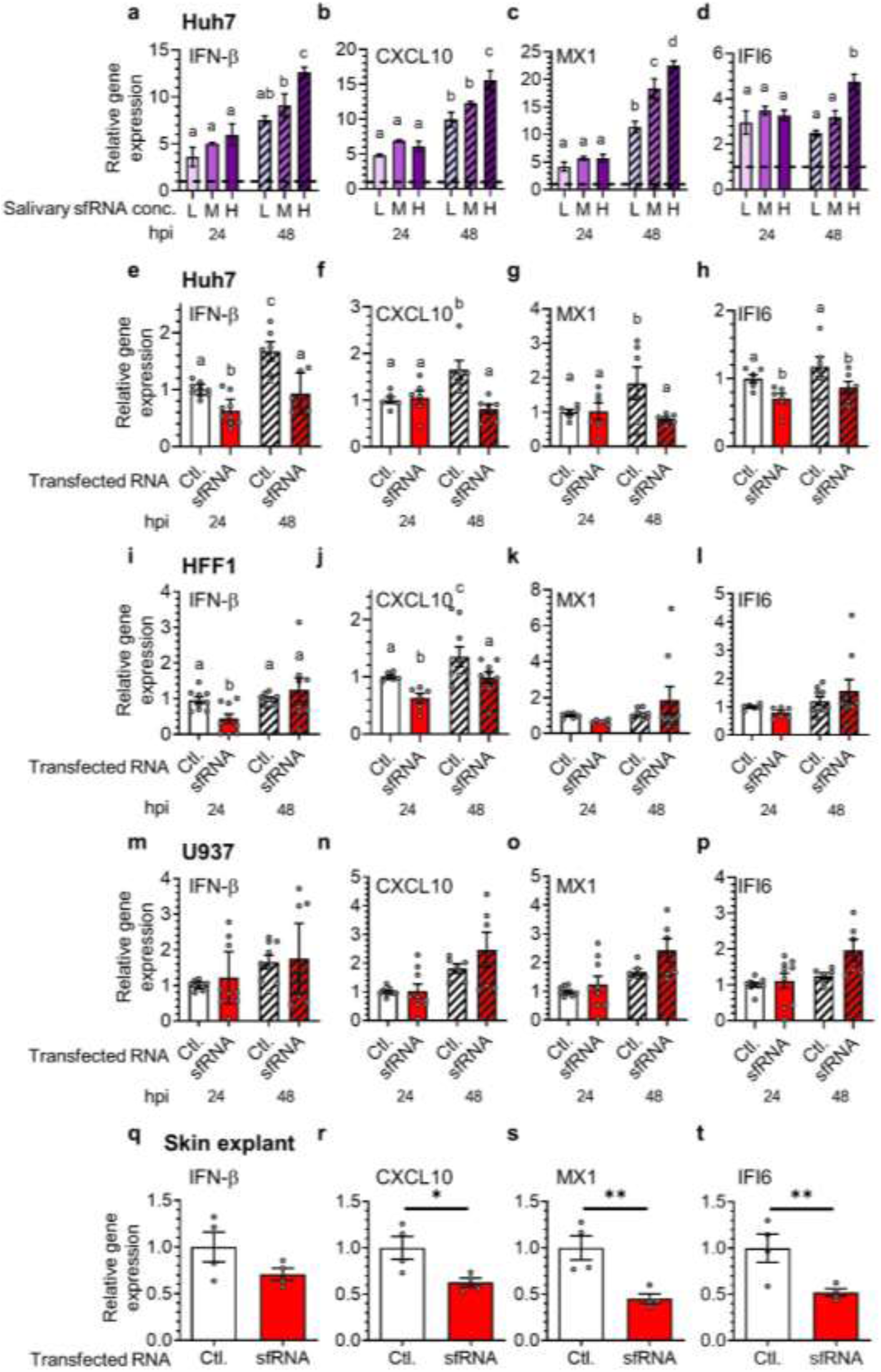
| Salivary sfRNA inhibits IFN responses. **a-d** Expression of IFN-β (a), CXCL10 (b), MX1 (c), and IFI6 (d) in Huh7 cells at 24 and 48 h post infection (hpi) with WNV- infected *Culex* saliva containing different concentrations of sfRNA. Category of sfRNA concentration; L, low; M, moderate; and H, high. N per category, 2. The dotted line indicates the mock-infected values. **e-h** Expression of IFN-β (e), CXCL10 (f), MX1 (g), and IFI6 (h) in Huh7 cells at 24 and 48 hpi with WNV at MOI 0.005 post sfRNA transfection. **i-l** Expression of IFN-β (i), CXCL10 (j), MX1 (k), and IFI6 (l) in HFF1 cells at 24 and 48 hpi with WNV at MOI 0.005 post sfRNA transfection. **m-p** Expression of IFN-β (m), CXCL10 (n), MX1 (o), and IFI6 (p) in U937 cells at 24 and 48 hpi with WNV at MOI 0.005 post sfRNA transfection. **q-t** Expression of IFN-β (q), CXCL10 (r), MX1 (s), and IFI6 (t) in skin explants at 24 hpi with WNV post sfRNA injection. Ctl., RNA control. Bars show mean ± sem. Repeats are represented by dots. Different letters show significant differences according to post hoc Fisher’s LSD test or T-test.

To determine whether sfRNA inhibits the innate immune response, we quantified the expression of the IFN-related genes in hepatocyte Huh7, fibroblast HFF1 and monocyte U937 cells that were transfected with sfRNA or the control RNA prior to WNV infection at MOI 0.005 and 0.5 (same samples as in Fig. 3h-j and S17a-c Fig.).

In Huh7 cells, sfRNA transfection inhibited the expression of *IFN-β* and *IFI6* at 24 and 48 hpi for both inocula, while *CXCL10* and *MX1* expressions were diminished at 48 hpi with the low inoculum only (Fig. 4e-h; S17d-g Fig.). In HFF1 cells, under low MOI infection, sfRNA strongly suppressed *IFN-β* expression at 24 hpi and reduced *CXCL10* expression at both 24 and 48 hpi (Fig. 4i-l). Under high MOI conditions, all four markers of IFN response were downregulated at 48 hpi (S17h-k Fig). In U937 cells, sfRNA transfection under low MOI did not inhibit the expression of the IFN-related genes (Fig. 4m-p); however, all four IFN markers were downregulated at 48 hpi, with *IFN-β* also reduced at 24 hpi under high MOI conditions (S17l-o Fig.). The lack of inhibition of certain IFN-related genes may reflect differences in the kinetics or intensity of gene expression among the different cell types.

Finally, we quantified the expression of IFN-related genes in human skin explants co-injected with WNV and either sfRNA or control RNA (same samples as in Fig. 3k). SfRNA moderately reduced *IFN-β* expression and significantly diminished *CXCL10*, *MX1* and *IFI6* expressions at 24 hpi (Fig. 4q-t). Together, these results demonstrate that salivary sfRNA dampens innate immune activation in the skin.

### SfRNA alters MDA5 signaling of interferon

To determine whether the early presence of sfRNA increases infection by altering the IFN response, we chemically inhibited IFN induction using the TBK1/IKKε inhibitor MRT67307, and infected Huh7 cells with WNV following transfection with either sfRNA or control RNA (Fig. 5a). Successful inhibition of the IFN response was confirmed by the lack of induction for *IFN-β*, *CXCL10*, *MX1* and *IFI6* expressions following control RNA transfection and infection (S20a-d Fig.). Supporting the hypothesis that sfRNA enhances infection by altering the IFN response, we observed that inhibition of IFN induction abolished sfRNA-mediated enhancement of infection at 24 and 48 hpi (Fig. 5b). The main pattern recognition receptors triggered by orthoflavivirus infection are RIG-I, MDA5 and TLR3 ^55^. Since Huh7 cells used here do not express TLR3 ^56^, we tested the role of RIG-I by using RIG-I-deficient Huh7.5 cells ^57^ (Fig. 5a). In Huh7.5 cells, the sfRNA-mediated enhancement of WNV infection was maintained at 24 and 48 hpi (Fig. 5c). Next, we silenced MDA5 in Huh7.5 to avoid any confounding effect from RIG-I (S21a,b Fig.). Strikingly, while sfRNA-mediated enhancement of infection was observed following control siRNA transfection, the infection enhancement was abolished at 24 and 48 hpi when MDA5 was silenced (Fig. 5d). Together, these findings indicate that early presence of sfRNA promotes infection by altering MDA5 downstream signaling.

**Fig. 5.**
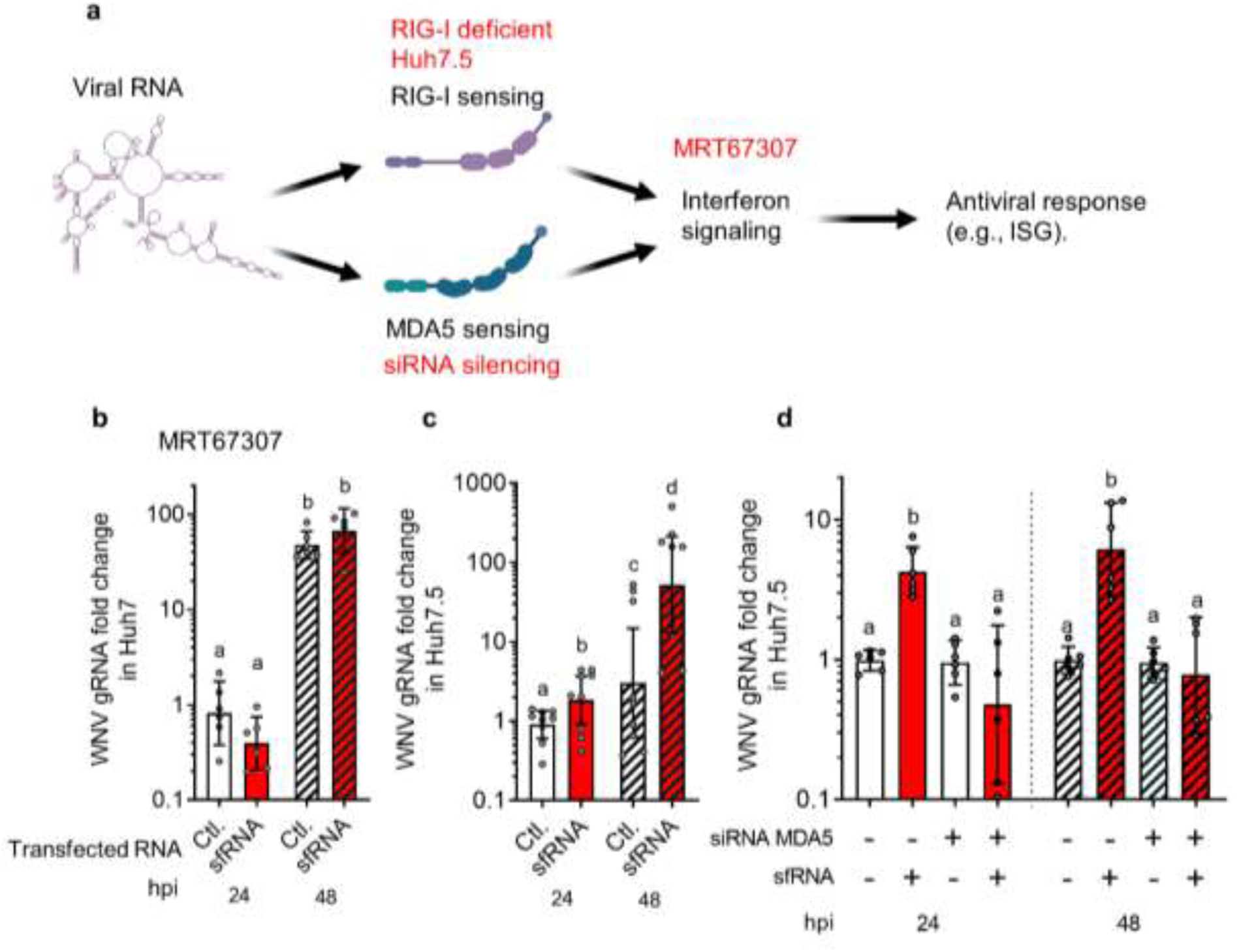
| sfRNA alters MDA-5 antiviral interferon response. **a** Scheme of RIG-I and MDA5 mediated antiviral interferon response. The methods used to disrupt the antiviral response are written in red. **b-d** WNV gRNA fold change post sfRNA transfection at 24 and 48 hpi in Huh7 cells treated with MRT67307 (b), in RIG-I-deficient Huh7.5 cells (c), and in Huh7.5 cells upon MDA5 silencing (d). Ctl., RNA control. Bars show geometric mean ± 95% C.I. Repeats are indicated by dots. Different letters show significant differences according to post hoc Fisher’s LSD test.

## Discussion

The incomplete understanding of orthoflavivirus transmission by mosquitoes hampers the identification of pan-orthoflavivirus targets that could be harnessed for broad-spectrum strategies to simultaneously protect against the multiple orthoflavivirus-related health threats. In this study, we identified and characterized sfRNA in mosquito saliva as a transmission-enhancing factor conserved across orthoflaviviruses. Using two phylogenetically distant orthoflaviviruses (WNV and ZIKV), belonging to distinct phylogenetic groups ^58^ and vectored by different genera of mosquitoes (*Culex* and *Aedes* mosquitoes), we showed that both orthoflaviviruses secrete sfRNA in salivary lipid particles. Combined with our previous discovery of DENV sfRNA in *Aedes* salivary vesicles ^23^ and our mechanistic characterization of sfRNA loading into salivary exosomes ^27^, the current study implies that sfRNA secretion in mosquito saliva is conserved across multiple, if not all, orthoflavivirus-mosquito species systems. We then designed a robust experimental approach to assess the function of salivary sfRNA in infection by infecting human cells and human skin explants with infectious mosquito saliva. This unprecedented dataset revealed that higher salivary sfRNA concentrations correlate with enhanced infection in human skin. We further demonstrated that sfRNA is responsible for the infection enhancement by reproducing this effect through the supplementation of *in vitro*-transcribed sfRNA in transmission-relevant cell types and human skin explants. To extrapolate from the *in vitro* and *ex vivo* findings and provide *in vivo* evidence for the role of salivary sfRNA in transmission, we used a mouse model of transmission. We established that sfRNA co-delivered with orthoflavivirus in the skin increases transmission and aggravates disease severity. Finally, we elucidated the mechanism by which the early presence of sfRNA enhances infection, demonstrating that sfRNA suppresses the early innate immune response and increases WNV infection by altering MDA5-mediated IFN induction. Altogether, our study establishes that multiple orthoflaviviruses secrete sfRNA in mosquito saliva to promote bite-initiated skin infection by inhibiting the innate immune response, thereby enhancing transmission.

Our results show that salivary sfRNA is shielded from nucleases by a detergent-sensitive layer, indicating the presence of a protective lipid membrane. These detergent-sensitive compartments could be either virions enclosed by a lipid bilayer envelop or extracellular vesicles (EVs). EVs are non-replicative, cell-derived membranous particles secreted into the extracellular space by most cells ^58^. We previously detected EVs in saliva of *Aedes* mosquitoes ^23^, while others observed EVs from mosquito cells ^59,60^. Multiple lines of evidence support that sfRNA is secreted inside EVs rather than virions, which by definition contain gRNA. Firstly, orthoflavivirus virions are spherical structures with a diameter of approximately 55 nm, which is too small to accommodate both gRNA and sfRNA ^61–63^. Secondly, we observed a divergence between sfRNA and gRNA quantities in SG and saliva, suggesting that sfRNA and gRNA produced in SG are not secreted through the same pathway. Thirdly, we showed that the particles containing sfRNA and gRNA differ in their sensitivity to nuclease degradation. Fourthly, the cytosolic origin of the EV lumen ^58,64^ would allow the loading of sfRNA. Indeed, sfRNA biogenesis, which results from gRNA degradation, occurs in the cytosol within processing bodies (PB) ^65^, a process supported in mosquitoes by the interaction of sfRNA with PB proteins such as Staufen ^66,67^. In contrast, virions are assembled in the ER and secreted via the trans-Golgi network ^7^. Overall, several viruses exploit EVs to secrete diverse types of viral RNA fragments ^68–71^ and our integrated datasets with DENV, WNV and ZIKV in *Aedes* and *Culex* mosquitoes suggest that orthoflaviviruses load sfRNA into mosquito EVs for salivary secretion. A study we published during the revision further support the secretion of sfRNA inside salivary exosomes ^27^.

We observed that salivary sfRNA favors bite-initiated skin infection by mitigating the early innate immune response. During the initial stages of orthoflavivirus infection, the innate immune system mounts a potent antiviral response mediated by IFN and involving ISGs ^15^. RIG-I and MDA5 are the two main viral RNA recognition receptors responsible for IFN activation via the IRF3/IRF7 phosphorylation signaling cascade ^72^. Loss-of-function studies for RIG-I and MDA5 have shown increased infection for WNV, ZIKV and DENV, establishing RIG-I and MDA5-mediated responses as major barriers to orthoflavivirus infection ^73–77^. We demonstrated that the infection enhancement by the early presence of WNV sfRNA was abrogated when MDA5 signaling was silenced, whereas RIG-I elimination did not affect sfRNA’s impact on infection. Our results indicate that salivary WNV sfRNA dampens the IFN response by mitigating MDA5 signaling in Huh7 cells, although the molecular target of sfRNA may vary across cell types. Consistent with sfRNA altering IFN upstream of IRF3/IRF7, one previous study reported that WNV sfRNA modulates IFN activation upstream of IRF3/IRF7 in murine embryonic fibroblasts (MEF) ^78^ and our findings show that chemical inhibition of IRF3/IRF7 activation abrogated the sfRNA effect. Alternatively, ZIKV sfRNA inhibits the IFN response by stabilizing NS5 inhibition of STAT1 phosphorylation – a step downstream of IRF3/IRF7 ^79^. DENV sfRNA inhibits RIG-I activation by binding TRIM25 to prevent RIG-I ubiquitylation ^26^ and interacts with G3BP1, G3BP2 and Caprin1 to reduce ISG translation ^80^. Despite differences in mechanisms, sfRNA from multiple orthoflaviviruses share the ability to inhibit the innate immune response ^24^, pointing to adaptive convergence in flavivirus evolution. However, our mechanistic contribution to WNV sfRNA function differs from previous studies as we evaluated the early presence of sfRNA prior to infection establishment and the production of other viral IFN inhibitors^7^.

Salivary sfRNA emerges as the first transmission-enhancing strategy linked to an orthoflaviviral component. Our study provides evidence that co-delivery of sfRNA at the bite site increases skin infection and exacerbates disease severity. Previous studies have underscored the importance of skin infection in transmission ^10,14^, and our findings showing the potent immune inhibition of sfRNA in fibroblasts - the primary dermal cell types - emphasize the critical role of orthoflavivirus replication in stromal cells before viral migration through myeloid cells ^81,82^. Based on our data and those of others, we propose a model in which the early presence of sfRNA in saliva prevents the initial activation of innate immune response to facilitate the establishment of skin infection and amplify subsequent viral dissemination. Our findings suggest that this model is valid for mosquito-borne orthoflaviviruses.

### Limitations of the study

While we propose that sfRNA secretion in salivary vesicles is a shared characteristic among orthoflaviviruses, our evidence is derived from the examination of only three orthoflavivirus species vectored by two distinct mosquito genera. Given that there are over 70 known orthoflavivirus species ^83^, including other highly pathogenic ones transmitted by mosquito genera not investigated here - such as for yellow fever virus, which is transmitted by *Haemagogus* mosquitoes ^84^ - our findings may not universally apply to all mosquito-borne orthoflaviviruses. Although sfRNA biogenesis and its anti-immune functions are conserved across all orthoflavivirus groups tested to date ^24,85,86^, it remains to be determined whether salivary sfRNA secretion contributes to transmission enhancement for all mosquito-borne orthoflaviviruses. Additionally, our characterization of the lipidic vesicles containing sfRNA is limited. The small volume of mosquito saliva restricts the extent of biochemical and biophysical analyses typically performed on EVs ^87^. Moreover, isolating sfRNA-containing vesicles from virions presents challenges due to the physical and chemical similarities between virions and lipidic vesicles ^70^.

## Materials and methods

### Cells and viruses

*Aedes albopictus* C6/36 and Human hepatocyte Huh7 cells were obtained from Cell Lines Services GmbH and Huh7.5 cells were kindly provided by Paul D. Biensiasz (The Rockefeller University, New York, NY, USA). Vero (CCL-81), HFF1 (SCRC-1041) and U937 (CRL-1593.2) cells were obtained from ATCC. Mammalian cell lines were grown in Dulbecco’s Modified Eagle Medium (DMEM) (Gibco), supplemented with 5 % heat-inactivated fetal bovine serum (FBS) (Eurobio) – except for HFF-1 which was grown with 15 % FBS - and 1 % penicillin/streptomycin mix (Invitrogen) at 37°C with 5 % CO2. Mosquito cells were grown in Roswell Park Memorial Institute (RPMI) media supplemented with 1 % non-essential amino-acids (ThermoFisher Scientific), 10 % FBS and 1 % penicillin/streptomycin mix at 28°C with 5% CO2.

WNV strain IS-98-ST1 (or Stork 98) was isolated from a stork in Israel in 1998 ^88^ and obtained from Dr. Philippe Desprès, Centre de Ressources Biologiques, Institut Pasteur, Paris. ZIKV strain H/PF13 was collected from human serum in French Polynesia in 2013 ^89^ and obtained from the European Virus Archive-Global (EVAg). Viruses were amplified in C6/36 cells, titrated in BHK21 cells as described ^40^ and stored in aliquots at -70°C.

### Mosquitoes

Experiments with *Culex* mosquitoes were carried out using the *Culex quinquefasciatus* SLAB strain, collected in California, USA ^90^, and obtained from the Institut des Sciences de l’Evolution in Montpellier, France. Experiments with *Aedes* mosquitoes were carried out using the *Aedes aegypti* Bora-Bora strain collected in French Polynesia in 1980 ^91^. Eggs from *Cx. quinquefasciatus* were hatched and reared at 25 ± 1°C, whereas eggs from *Ae. aegypti* were reared at 27 ± 1°C. Both species were reared at 70 ± 5% relative humidity and 12h:12h day:night. Larvae were distributed in plastic trays at a density of 200 individuals per tray and fed half a tablet of concentrated yeast and 1 g of TetraMin (Tetra) on the day of hatching, and then 1.5 g of TetraMin every two days until pupation. Pupae were transferred to cages and supplied with 10 % sugar solution and water *ad libitum*.

### Mice

C57BL/6J male mice were purchased from Charles-River (France) and housed in ventilated cages in NexGen Mouse 500 (Allentown; Serial number: 1304A0078) in the biosafety level 3 animal facility at MIVEGEC-IRD, Montpellier, France. Mice were maintained with a 17h:7h light/dark cycle, 53-57 % humidity, 20-24°C temperature and provided with an irradiation-sterilized mouse diet (A03, SAFE, France) and sterilized water *ad libitum*. Upon arrival, mice were let to rest for one week before experiments. Animal protocols were approved by the national ethical committee (permission numbers: 43466; 31273).

### Mosquito oral infection

Four-day-old female *Cx. quinquefasciatus* were starved for 12h, and offered an infectious blood meal for 60 min at sunset time, using the Hemotek feeding system (Discovery Workshops) with chicken skin (obtained from spring chicken purchased in supermarket). The artificial blood meal contained WNV at 8 × 10^5^ plaque forming unit (PFU)/ml, 50 % volume of washed erythrocytes from rabbit’s blood (animals housed in the BSL2 VectoPole animal facility, authorization number: H3417221), 25 mM ATP (ThermoFisher Scientific), 5 % FBS (Eurobio-scientific) and RPMI to complete the volume to 2.5 ml. Engorged females were selected and maintained in the rearing conditions for 10 days before analysis.

### Mosquito inoculation

Five-day-old cold-anesthetized female *Cx. quinquefasciatus* or *Ae. aegypti* were intrathoracically inoculated with 69 nl containing 100 PFU of WNV or 50 PFU of ZIKV, respectively, using Nanoject II (Drummond Scientific Company) and needles made from 1.14 mm O.D. glass capillaries x 1.75″ length (Drummond). Mosquitoes were then maintained in rearing conditions for 10 days before analysis.

### Mosquito salivary glands dissection and saliva collection

Cold-anesthetized mosquitoes had their wings and legs removed before inserting individual proboscises into 20 µl filter-sterile tips containing 10 µl of RPMI (for RNase resistance assay) or of DMEM (for human cell infection with WNV) with 2% 25 mM Erioglaucine (SigmaAldrich) for 30 min at 25°C. Mosquitoes with a blue abdomen (color of Erioglaucine) were considered to have salivated and the corresponding media were collected either individually or as pools. Salivary glands were dissected after salivation, labeled to identify the associated saliva and homogenized using Fast-Prep bead bitter homogenizer (MP) with glass beads.

### sfRNA production

Full length *in vitro*-transcribed sfRNAs from WNV and ZIKV were generated by amplifying the sfRNA sequence from virus cDNA using the following primer pair with T7-tagged forward: for WNV, 5’-TAATACGACTCACTATAGGGAGTCAGGCCGGGAAGTTCC-3’ and 5’-AGATCCTGTGTTCTCGCACC-3’; for ZIKV, 5’-TAATACGACTCACTATAGGGTCTTAATGTTGTCAGGCCTGCTA-3’ and 5’-CCGCTATTCGGCGATCTGT-3’. Amplicons were reverse transcribed using Megascript T7 kit (Ambion), monophosphorylated at the 5’ end using RNA 5’ Polyphosphatase (Lucigen), extracted in RNase-free water (Invitrogen) using E.Z.N.A. Total RNA kit I (Omega) to remove the polyphosphatase and folded by slowly reducing the temperature from 95 to 4°C with 5 mM of Mg^2+^, which regulates sfRNA folding ^92^. sfRNA quantity was estimated using nanodrop (ThermoFisher Scientific) and used to calculate copy number.

As control, a same-size RNA fragment (i.e., 483 nt) corresponding to a part of the dengue virus NS2 gene was *in vitro*-transcribed using the primers 5’-TAATACGACTCACTATAGGGGCAGCTGGACTACTCTTGAG-3’, 5’-GGTCCTGTCATGGGAATGTC-3’ ^66^, monophosphorylated and folded as for sfRNA.

### Northern Blot

Northern blot was conducted using NorthernMax Kit (Ambion) with modifications to manufacturer’s protocol as described ^23^. Total RNA from 100 infected mosquitoes or from saliva samples from about 300 infected mosquitoes was extracted using TRIzol reagent (Invitrogen) and separated on a denaturing gel with 5 % Acrylamide/Bisacrylamide 19:1 (Starlab) and 8 M Urea (Invitrogen). Biotinylated single-stranded RNA ladder (Kerafast) was also loaded on the gel. RNA was transferred onto a Hybond-N+ nylon membrane (Merck) using Trans-Blot Turbo (Bio-Rad) at constant 1.3 A for 30 min with Voltage ˂ 25 V. The membrane was UV-crosslinked, pre-hybridized and hybridized overnight with a biotin-16-dUTP (Roche) labeled dsDNA probe (1 mg) generated from the primers used to amplify the qPCR sfRNA targets for WNV and the primers used to generate full-length sfRNA for ZIKV. After washes, the membrane was blocked using Odyssey Blocking Buffer (LI-COR) and stained with IRDYE 800cw streptavidin (LI-COR) in TBS. Pictures were taken with ChemiDoc (Bio-Rad).

### Absolute quantification of gRNA copies

Total RNA was extracted using E.Z.N.A. Total RNA kit I (Omega Bio-Tek). gRNA for WNV was quantified using iTaq Universal SYBR Green one-step RT-qPCR kit (Bio-Rad) with primers: 5’-ATTCGGGAGGAGACGTGGTA-3’ and 5’-CAGCCGCCAACATCAACAAA-3’; in LighCycler 96 thermocycler (Roche) with the following thermal profile: 50°C for 10 min, 95°C for 2 min, and 40 cycles at 95°C for 15s, 60°C for 15s and 72°C for 20s. gRNA for ZIKV was quantified using iTaq Universal Probes one-step RT-qPCR kit (Bio-Rad) with primers : 5’-TTGGTCATGATACTGCTGATTGC-3’ and 5’-CCTTCCACAAAGTCCCTATTGC-3’ and probe: 5’-CGGCATACAGCATCAGGTGCATAGGAG-3’; in LighCycler 96 (Roche) with the following thermal profile: 50°C for 15 min, 95°C for 2 min, and 45 cycles at 95°C for 10s and 60°C for 30s.

Absolute quantification for both gRNAs was obtained by generating standard equations using *in vitro-*transcribed RNA qPCR targets. RNA targets for WNV and ZIKV were produced by amplifying virus cDNA with primer pairs: 5’-TAATACGACTCACTATAGGGATTCGGGAGGAGACGTGGTA-3’ / 5’-CAGCCGCCAACATCAACAAA-3’, and 5’-TAATACGACTCACTATAGTTGGTCATGATACTGCTGATTGC-3’ / 5’-CCTTCCACAAAGTCCCTATTGC-3’, respectively. Amplicons were reverse transcribed using Megascript T7 kit (Ambion), extracted using RNeasy Mini Kit (Qiagen) and quantified using nanodrop (ThermoFisher Scientific) to calculate copy number. Serial dilutions were used to establish absolute standard equations. Using the gRNA templates, the limit of detection (LoD) at 95% was determined by calculating fractions of detected samples in three replicates of serial dilutions ^93^.

gRNA detection rate was calculated as the proportion of samples with detectable amount of gRNA among all samples.

### Absolute quantification of sfRNA copies

sfRNA and 3’UTR were jointly quantified in total RNA extracts from E.Z.N.A. Total RNA kit I using the following primers for WNV: 5’-AGTTGAGTAGACGGTGCTGC-3’ and 5’-CCGTAGCGTGGTCTGACATT-3’, and for ZIKV: 5’-GCTGGGAAAGACCAGAGACT-3’ and 5’- CTATTCGGCGATCTGTGCCT-3’. Two different qPCR conditions were applied for each pair of primers. First, quantification for both viruses was conducted using iTaq Universal SYBR Green one-step RT-qPCR kit (Bio-Rad) in LightCycler with the following thermal profile: 50°C for 10 min, 95°C for 2 min, and 40 cycles at 95°C for 10s and 60°C for 25s. Second, quantification with WNV primers was conducted in the same conditions but for the reverse transcription step set at 60°C for 20 min. Additionally, quantification with ZIKV primers was conducted using a two-step RT-qPCR. RT was conducted with the reverse primer and M-MLV enzyme (Promega) with a denaturation/annealing step at 70°C and following manufacturer’s conditions with RT at 37°C. qPCR was then performed in AriaMx thermocycler (Agilent) using EvaGreen qPCR MixPlus (Euromedex) with the following thermal profile: 95°C for 15 min, and 40 cycles at 95°C for 15s, 60°C for 20s and 72°C for 20s.

Absolute quantification and LoD for sfRNA for WNV and ZIKV were obtained as described above for gRNA but using dilutions of sfRNA copies.

sfRNA copy number was calculated by subtracting the number of 3’UTR + sfRNA copies to the number of gRNA copies. For samples that contained detectable amount of gRNA, sfRNA:gRNA ratio was calculated by dividing the number of sfRNA over the number of gRNA. sfRNA detection rate was calculated as the number of samples with detectable amount of sfRNA only among gRNA-positive samples.

### Splinted-ligation reverse-transcription PCR (SL-RT-PCR)

RNA was mixed with 20 pmol of DNA splint oligo (5’-GCTGATGGCGATGAATGAACACTGCGTTTGCTGGCTTTGATGGAAAGTCAGGCCGGGAAGTTCCCGCCACCG-3’), 30 pmol of RNA anchor (5’-GCUGAUGGCGAUGAAUGAACACUGCGUUUGCUGGCUUUGAUG-3) in 8 µL total volume reaction using DEPC-treated water (ThermoFisher). Annealing was conducted at 95°C for 5 min, 70°C for 5 min, 60°C for 5 min, 42°C for 5 min, and 25°C for 5 min. Ligation was conducted with 200 U of T4 DNA ligase (NEB) in 1X T4 DNA ligase buffer and 20 U of RNase OUT (Thermo Scientific) at 16°C for 18 h. DNA digestion was performed by incubation with 5 U of RNase-free DNase I (Thermo Scientific) in 1X DNase buffer with MgCl2^+^ at 37°C for 3 h. RT-PCR amplification was performed using one-step RT-PCR (Bio-Rad) and 300 nM of RNA anchor-matching forward (5’-GCTGATGGCGATGAATGAACACTGC-3’) and sfRNA-matching reverse (5’-GCAGGCAGCACCGTCTACT-3’) primers in total volume of 10 µl in LightCycler 96 (Roche) with the following thermal profile: 50°C for 10 min, 95°C for 1 min, followed by 40 cycles of 95°C for 10 sec, 60°C for 25 sec. Amplicons were run on 3,5% agarose gel in 0.5X TAE buffer, stained with GelRed, and visualized under UV exposure (Vilber).

As control, a 5’ end fragment of WNV sfRNA (5’-GAAAGUCAGGCCGGGAAGUUCCCGCCACCGGAAGUUGAGUAGACGGUGCUG CCUGC-3’) synthetized by Eurofins (EU) was monophosphorylated with 5 U of T4 ploynucleotide kinase (NEB), 1X Buffer, 1 mM ATP in 25 µl total reaction at 37°C for 30 min and purified using EZNA Total RNA kit I (Omega).

### RNase resistance assay

10^7^ copies of *in vitro*-transcribed sfRNA or of viral RNA control were mixed with 10 saliva samples from uninfected mosquitoes and subjected to RNAse resistance assay. Pools of 40 saliva samples from intrathoracically-inoculated mosquitoes were divided into 4 equal volumes and subjected to the different conditions of the RNAse treatment assay. Volumes were adjusted to 100 µl with PBS. For WNV, samples were supplemented with 10 µl of 0.1 % Triton X-100 (Sigma) for 30 min at 4°C, before adding 0.5 µl of Micrococcal nuclease (MNase) (Thermo Scientific) and 10 µl of buffer (50mM Tris-HCl pH 8; 10mM CaCl2 final concentration) for 1h at 37°C. Controls for Triton X-100 and RNase were added the same volume of PBS instead and maintained at the corresponding temperatures. RNA was then extracted using QIAamp viral RNA kit (Qiagen).

For ZIKV, the same treatments were applied except that 3 µL of RNase A/T1 (ThermoFisher Scientific) was added for 30 min at 37°C instead of MNase.

### Virus titration of saliva

100,000 Vero cells per well of 24-well plates were incubated with 125 μl of 10-fold serial dilutions of saliva-containing media for 1h at 37°C with 5% CO2. After removing the inoculum, 1 ml of sterile 2% CarboxylMethyl Cellulose (CMC, SigmaAldrich) in DMEM supplemented with 2% FBS and 1% P/S was added. Four days later, cells were fixed with 4% paraformaldehyde (Sigma-Aldrich) for 20 min, covered with crystal violet solution (VWR) for 45 min, and rinsed. Plaques were counted using binocular microscope and plaque forming unit (PFU) per ml was calculated using the formula: number of plaques x 8 x dilution factor.

gRNA:PFU ratio was calculated by dividing the gRNA copy number quantified as detailed above to the PFU number within the same saliva.

### Infection of human cells with WNV infectious *Culex* saliva

Pools of 10 saliva samples from *Culex* mosquitoes infected by oral feeding with 10^8^ PFU/ml of WNV were collected and stored at -70°C. gRNA and sfRNA were quantified from a frozen aliquot of each pool. 5,000 Huh7 cells per well of a 96-well flat-bottomed plate were inoculated with 1,000 gRNA copies of the different saliva pools for 2h. Total volume of the infected saliva was homogenized to 50 µl by adding uninfected saliva from *Culex* female mosquitoes. After infection, inoculum was removed and cells were supplemented with 2 % FBS DMEM and collected at 24 and 48h post inoculation by adding TRK lysis buffer and total RNA was extracted using E.Z.N.A. total RNA kit I.

### Injection of WNV infectious *Culex* saliva into human skin explants

Five mm diameter Human skin explants obtained from Biopredic International (France) were decontaminated by washing in 25 mM HEPES (Gibco), 400 U/ml P/S and 10 µg/ml amphotericin B (biowest) in DMEM. Explants were maintained individually in 8 µm cell culture insert (Thincert) with 200 µl of two-week skin culture medium w/o animal components (MIL215C, Biopredic) at 37°C, 5% CO2. Explants were inoculated with 760 gRNA copies of saliva pools. Total volume of the infected saliva was homogenized to 30 µl by adding uninfected saliva from *Culex* female mosquitoes. At 24 h post injection, explants were cut into small pieces and, homogenized using the bead bitter homogenizer and glass beads before RNA extraction as detailed for mouse skin above.

### Relative quantification of WNV gRNA and IFN-related genes in human cells and skin explants

RNA was reverse transcribed using the PrimeScript RT Reagent Kit (Perfect RealTime, Takara Bio Inc.). Real-time PCR reaction was performed in duplicate using Takyon ROX SYBR MasterMix blue dTTP (Eurogentec) on an Applied Biosystems QuantStudio 5 (Thermo Fisher Scientific) in 384-well plates. Transcripts were quantified using the following program: 3 min at 95°C followed by 35 cycles of 15 s at 95°C, 20s at 60°C, and 20s at 72°C. Values for each transcript were normalized to the geometric means of Ct values of 4 different housekeeping genes (*RPL13A*, *ACTB*, *B2M*, and *GAPDH*) using the 2^-ΔΔCt^ method. Primers used for quantification of transcripts by real-time qPCR were: RPL13A 5’-AACAGCTCATGAGGCTACGG-3’ and 5’-TGGGTCTTGAGGACCTCTGT-3’, ACTB 5’-CTGGAACGGTGAAGGTGACA-3’ and 5’-AAGGGACTTCCTGTAACAATGCA-3’, B2M 5’-TGCTGTCTCCATGTTTGATGTATCT-3’ and 5’-TCTCTGCTCCCCACCTCTAAGT-3’, GAPDH 5’-TGCACCACCAACTGCTTAGC-3’ and 5’-GGCATGGACTGTGGTCATGAG-3’, IFN-β 5’-TGCTCTCCTGTTGTGCTTCTC-3’ and 5’-CAAGCCTCCCATTCAATTGCC-3’, CXCL10 5’-CGCTGTACCTGCATCAGCAT-3’ and 5’-GCAATGATCTCAACACGTGGAC-3’, Mx1 5’-AAGCTGATCCGCCTCCACTT-3’ and 5’-TGCAATGCACCCCTGTATACC-3’, IFI6 5’-GGGTGGAGGCAGGTAAGAAA-3’ and 5’-GACGGCCATGAAGGTCAGG-3’. Primers for WNV gRNA were as described above.

### Infection of human cells post sfRNA transfection

2 × 10^5^ Huh7, HFF1, U937 or Huh7.5 cells were transfected with 10^10^ copies of either monophosphorylated-folded-sfRNA or the control RNA using TransIT-mRNA Transfection Kit (Mirus) for 2 h. The transfection media was removed and cells were immediately infected with WNV at a multiplicity of infection, MOI, of 0.005 or 0.5 for 1h. Cells were then maintained in 2% FBS media for 24 and 48 h before extracting RNA using E.Z.N.A. total RNA kit I.

### Co-injection of sfRNA and WNV in human skin explants

Five mm diameter human skin explants as detailed above were inoculated with 10^2^ PFU of WNV mixed with 10^9^ copies of monophosphorylated-folded-sfRNA or the control RNA in a total volume of 4 µl using a Nanoject II injector (Drummond) and glass needles. At 24 h post injection, explants were cut into small pieces and, homogenized using the bead bitter homogenizer and glass beads before RNA extraction as detailed for mouse skin above.

### Co-injection of sfRNA and WNV in mice

Five-to-seven-week-old mice were shaved with the animal trimmer (VITIVA MINI, BIOSEB) on the lower back one day prior to injection to limit shaving-induced inflammation. Mice anesthetized by intraperitoneal injection of 0.2 ml/mouse of 10 mg/ml of ketamine (Imalgène 1000, Boehringer Ingelheim Animal Health) and 1 mg/ml of xylazine (Rompun 2%, Elanco GmbH) were intradermally inoculated with 10^3^ PFU of WNV mixed with 10^10^ copies of monophosphorylated-folded-sfRNA or the same number of the control RNA. The total volume of inoculum was 4 µl. As an additional control, other mice were only injected with monophosphorylated-folded-sfRNA.

At 24 h post-injection, skin biopsies from the injection site and draining lymph nodes were collected and homogenized in 340 µL of TRK lysis buffer using Fast-Prep bead bitter homogenizer (MP) set at 1.4 m/s for 60 sec. with 1 nm glass beads. Total RNA was extracted using E.Z.N.A. total RNA kit I. At 4 days post injection, blood samples were collected via mandibular puncture. RNA was extracted using QIamp viral RNA kit (Qiagen) and WNV gRNA per ml of blood was absolutely quantified after evaluating blood volume by pipetting. Clinical signs and weight were registered daily to determine a clinical score (CS) and calculate the percentage of weight loss until 14 days post-injection. CS of 0 was assigned to healthy mice; 1 for mice with ruffled fur, lethargy, hunched posture, no paresis, normal gait; 2 for mice with altered gait, limited movement in 1 hind limb; 3 for lack of movement, paralysis in 1 or both hind limbs; 4 for moribund mice; and 5 for dead mice ^94^. Mice were euthanized under anesthesia if they displayed neurological symptoms, severe distress, or weight loss exceeding 20%, or reached 12 days post-injection.

### Chemical inhibition of interferon signaling

Media of 2 × 10^5^ Huh7 cells was supplemented with 10 µM of MRT67307 (MedChemExpress) diluted in water for 1h, then during transfection of monophosphorylated-folded-sfRNA or the control RNA, and infection with WNV as described above.

### MDA-5 Silencing

2 × 10^5^ Huh7.5 cells were transfected overnight with 5 pmol of multiplex siRNAs (SI05130608, Qiagen) using Lipofectamine RNAiMax (ThermoFisherScientific). As a control, Allstars multiplex siRNAs (SI91027281, Qiagen) were similarly transfected. 48 h later, cells were transfected with monophosphorylated-folded-sfRNA or the RNA control and subsequently infected with WNV as described above.

### Western blot

2 × 10^5^ Huh7.5 cells were lysed with 100 µl of RIPA (Biotech) supplemented with protease inhibitor cocktail (cOmplete, Roche). After centrifugation at 14,000 g for 15 min at 4°C, proteins were quantified in the supernatant using Qubit protein assay (ThermoFisherScientific). Normalized quantities of protein extracts were separated on NuPAGE 4-12% acrylamide gel (Invitrogen) and transferred onto 0.2 µm Nitrocellulose membrane using a Trans-Blot Turbo Transfer System (Bio-Rad). Membranes were blocked with 5% skimmed milk (ThermoScientific), incubated with 1:4,000 anti-MDA5 antibody (21775-1-AP, Proteintech) or 1:400 anti-Actin (MA5-11869, Invitrogen) in blocking solution, washed with 0.05% tween-20 PBS, incubated with 1:500 anti-rabbit IgG HRP-linked antibody (Cell signaling) in blocking solution, and washed. Blots were revealed using SuperSignal West Pico PLUS (ThermoScientific) and observed with ChemiDoc Imaging System (BioRad).

### Statistical analysis

Comparison of gRNA and sfRNA copies, relative gRNA fold changes and IFN-related genes were conducted using one tailed T-test or post-hoc Fisher’s LSD test. Absolute number and relative values of gRNA and sfRNA copies were log-transformed before statistical analysis. Correlation analysis was performed using linear regression analysis. Survival analysis were performed with Gehan-Breslow-Wilcoxon test. The statistical analyses were performed using GraphPad Prism v9.0.

The progression of clinical scores in mice was analyzed using a general linear mixed model with a negative binomial distribution to account for overdispersed data. The progression of weight change in mice was analyzed using a general linear mixed model with an inverse Gaussian distribution and log link function, appropriate for non-negative, right-skewed data. The model assessed the effects of days post-infection and experimental groups (WNV + sfRNA vs. WNV + Ctl.), including random intercepts for individual mice to account for variability across subjects. For pairwise comparisons between groups, post hoc analyses were conducted using estimated marginal means (emmeans) with Tukey’s adjustment for multiple comparisons. All analyses were conducted in R (version 4.3.3) using the lme4 ^95^, glmmTMB ^96^, and emmeans packages ^97^.

## Data availability

All data supporting the findings of this study are available within the paper and its Supplementary Information.

## Acknowledgments

We thank the VectoPole in Montpellier and Arnaud Berthomieu for providing mosquito eggs. We also thank Audrey Vernet from the proteomic platform. PhD scholarships for IS was provided by the Institut Méditerranéen Hospitalier (IHU, Marseille), and for FRC and FR by the French Ministry of Research and Higher Education (CBS2 doctoral school, Montpellier). Post-doctoral fellowships from Fondation pour la Recherche Médicale to HM (SPF202110013925) and to EFM (ARF202309017577). Support for the research was provided by the French Agence Nationale pour la Recherche (ANR-20-CE15-0006) and by the EU (HORIZON-HLTH-2023-DISEASE-03 #101137006) to JP.

## Competing interests

The authors declare no competing interests.

## Author contributions

Conceptualization: ISP, JP Formal analysis: ISP, JP Funding acquisition: JP

Investigation: ISP, HM, LB, KJ, LP, NS, MT, JZ, FR, ZR, QN, FRC, SR, SF, CM, CG, BS, WS, EM

Methodology: ISP, HM, AG, DRS, RH, OM, DM, SN, JP

Project administration: JP Supervision: OM, JP Visualization: ISP, JP

Writing – original draft: ISP, JP Writing – review & editing: all authors

## Notes

### Competing Interest Statement

The authors have declared no competing interest.

### Summary of Updates

This revised version include the experiments and information requested during the review process.

