## Supplemental material for "Multiple orthoflaviviruses secrete sfRNA in mosquito saliva to promote transmission by inhibiting MDA5-mediated early interferon response"

### Supplementary material

#### Results

##### Optimization of WNV sfRNA quantification

Quantification of sfRNA using a one-step RT-qPCR approach, specifically targeting sfRNA1, initially failed to detect sfRNA (data not shown). To quantify sfRNA, we employed an indirect subtractive approach. We quantified the combined copy number of sfRNA and 3'UTR with primers that anneal onto both sfRNA and 3'UTR. We then subtracted the number of gRNA to calculate sfRNA copies, as previously implemented for DENV [24,48]. The calculation of sfRNA+3'UTR copies is based on a standard equation generated from known amounts of *in vitro*-transcribed full-length sfRNA target, which may not fold as *in cellulo*-produced sfRNA. We reasoned that the RT-qPCR protocol had a reduced efficiency in amplifying structured sfRNA [49], resulting in an undervaluation of *in cellulo*-produced folded sfRNA. Accordingly, we observed that the RT-qPCR protocol with reverse transcription at 50°C was less efficient in amplifying structured than unfolded *in vitro*-transcribed sfRNA (S2a,b Fig.). We therefore adjusted the reverse transcription temperature to 60°C to unfold sfRNA and observed a similar amplification efficiency for both folded and unfolded sfRNA (S2c,d Fig.), ensuring accurate sfRNA quantification.

##### Optimization of ZIKV sfRNA quantification

Similar to the challenges encountered with WNV sfRNA, quantification of ZIKV sfRNA using one-step RT-qPCR with reverse transcription at 50°C proved less efficient for folded than unfolded sfRNA (S6a,b Fig.). To overcome this, we optimized the quantification process by developing a two-step RT-qPCR with a denaturation/annealing step at 70°C, in order to unfold sfRNA structures and detect unfolded and folded sfRNA equally well (S6c,d Fig.).

#### Supplementary figures

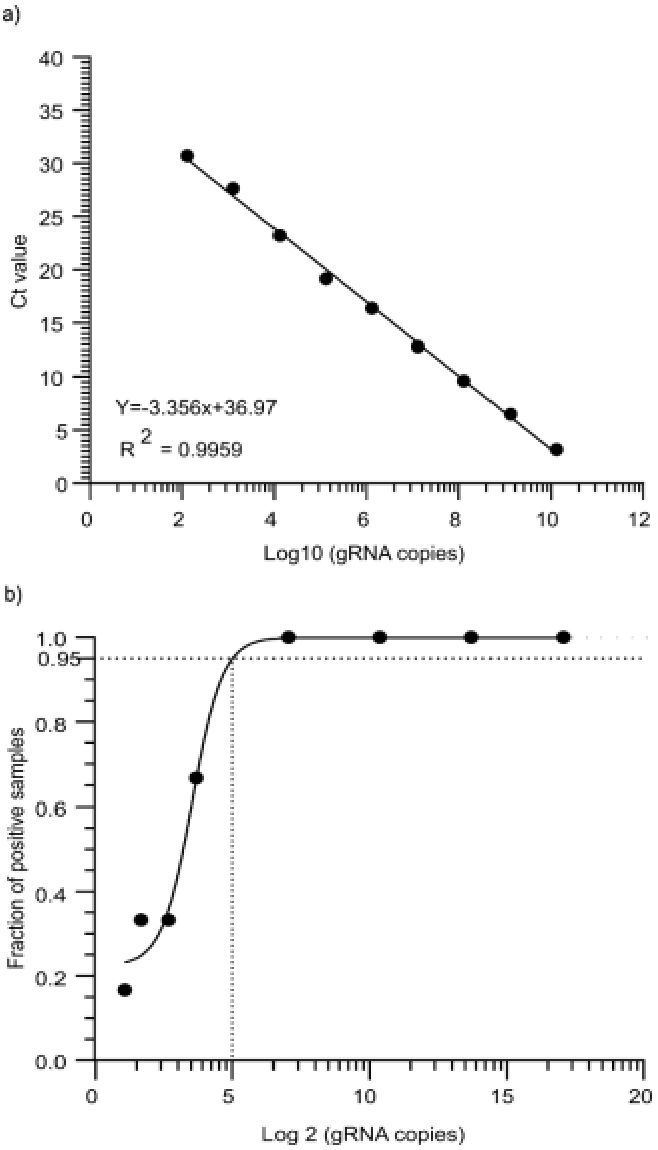

**S1 Fig. (Related to Fig. 1) | Absolute quantification of WNV gRNA.**

In vitro transcribed RT-qPCR target was serially diluted from 1.29 x 1010 to 2 copies, before quantification by one-step RT-qPCR. a Standard curve for absolute quantification of gRNA copies. Points show averaged Ct calculated from three independent replicates. b Limit of Detection (LoD) at 95% confidence for WNV gRNA is 25 copies ($5^2$). Fractions of positive samples were calculated from six replicates.

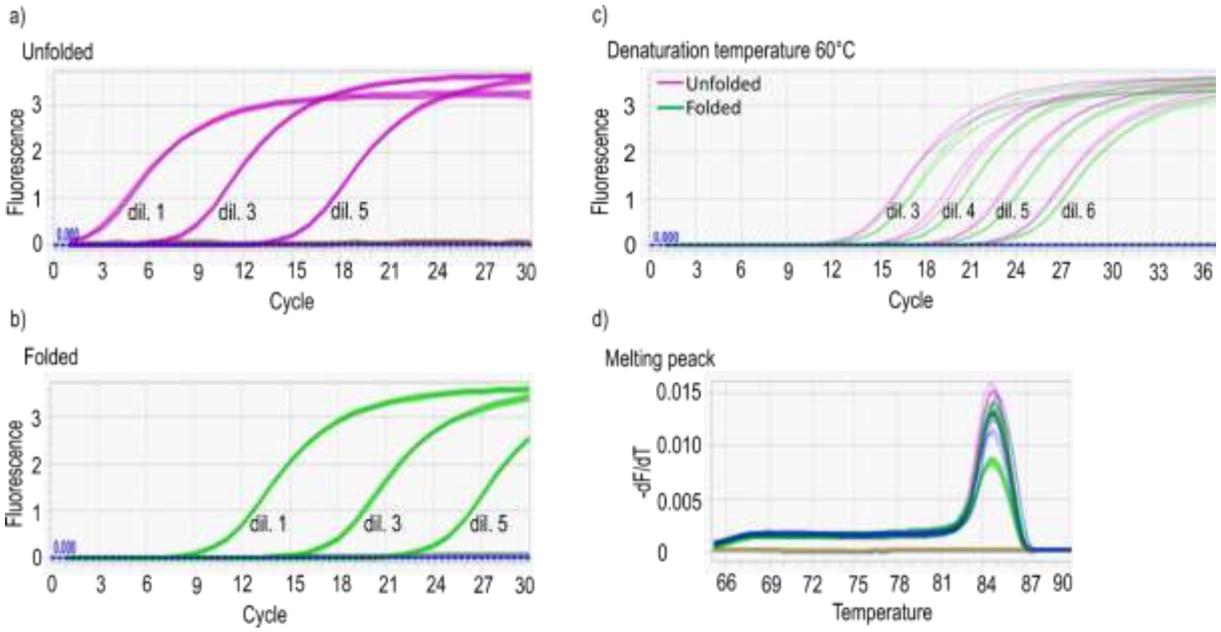

**S2 Fig. (Related to Fig. 1) | Optimization of RT-qPCR quantification for folded WNV sfRNA.**

a-b Amplification curves for different dilutions (dil. 1, 3 and 5) of unfolded (a) and folded (b) *in vitro*-transcribed WNV sfRNA in one-step RT-qPCR with RT temperature at 50°C. c Amplification curves for the same dilutions (dil. 3-6) of unfolded and folded *in vitro*-transcribed WNV sfRNA in one-step RT-qPCR with RT temperature at 60°C. d Melting curve analysis for the reaction in c.

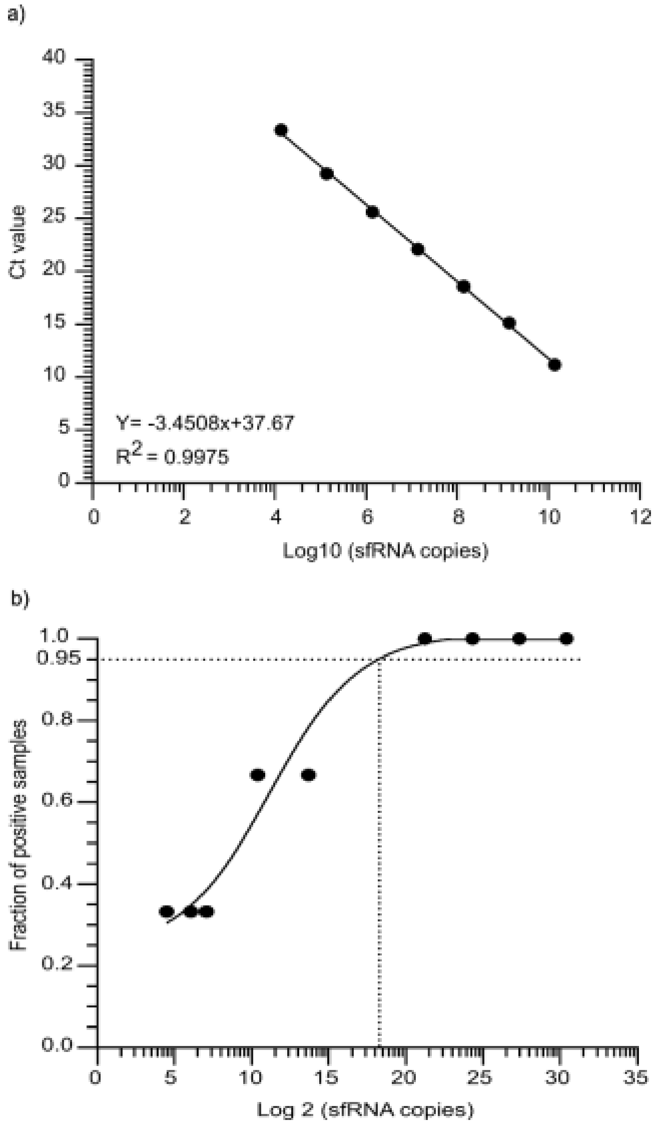

**S3 Fig. (Related to Fig. 1) | Absolute quantification of WNV sfRNA**.

*In vitro* transcribed RT-qPCR target was serially diluted from 1.38 x $10^{10}$ to 22 copies, before quantification by one-step RT-qPCR with RT at 60°C. **a** Standard curve for absolute quantification of sfRNA copies. Points show averaged Ct calculated from three independent replicates. **b** The limit of Detection (LoD) at 95% confidence for WNV sfRNA is 342 copies ($18.5^2$). Fractions of positive samples were calculated from tree replicates.

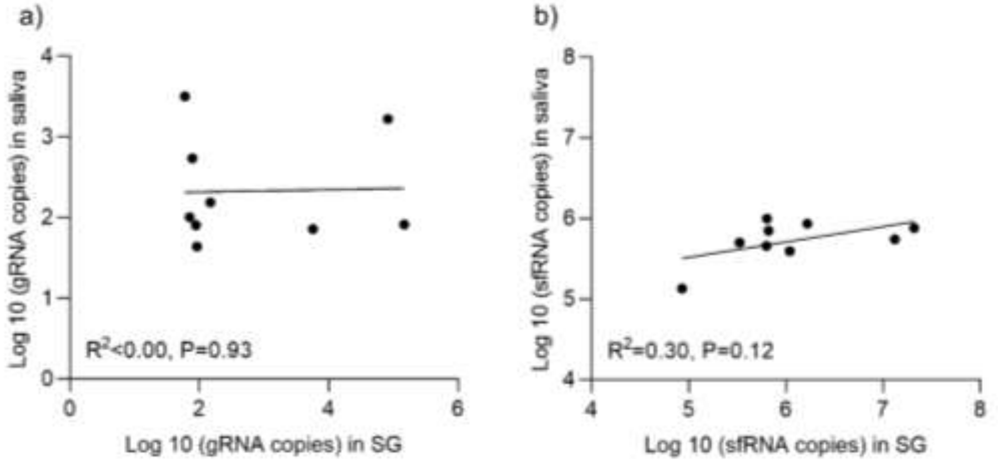

**S4 Fig. (Related to Fig. 1) | a-b Correlations between salivary glands (SG) and saliva for gRNA and sfRNA copies for WNV orally infected mosquitoes**.

Samples were collected from *Culex* mosquitoes orally infected with 8 x $10^5$ pfu/ml. Dots indicate SG and saliva repeats. P values were determined by correlation analysis.

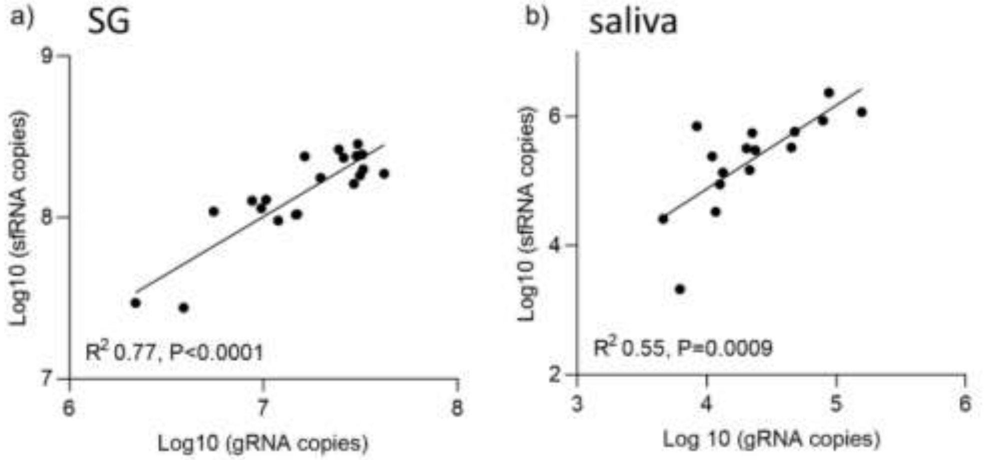

**S5 Fig. (Related to Fig. 1) | a-b Correlations between gRNA and sfRNA copies in SG and saliva from WNV-inoculated mosquitoes.**

Dots indicate SG and saliva repeats. P values were determined by correlation analysis.

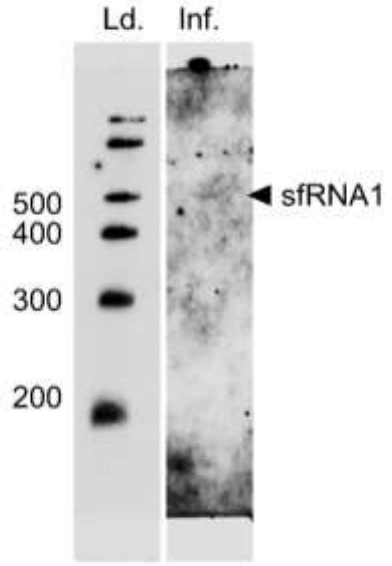

**S6 Fig. (Related to Fig. 1) | Northern blot detection of sfRNA in saliva from WNV-infected mosquitoes.**

Detection of WNV sfRNA in inoculated *Culex* mosquitoes. Ld., RNA ladder; Inf., WNV-infected *Aedes* mosquitoes.

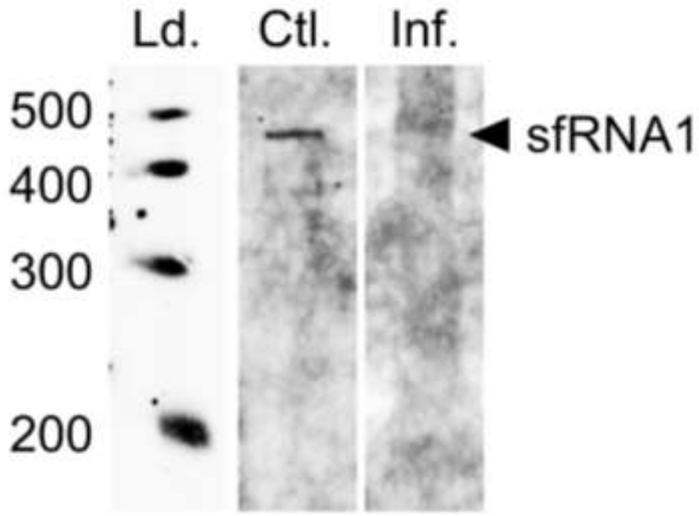

**S7 Fig. (Related to Fig. 1) | Northern blot detection of ZIKV sfRNA in whole mosquitoes.**

Detection of ZIKV sfRNA in inoculated *Aedes* mosquitoes. Ld., RNA ladder; Ctrl., *in vitro* transcribed folded ZIKV sfRNA1; Inf., ZIKV-infected *Aedes* mosquitoes.

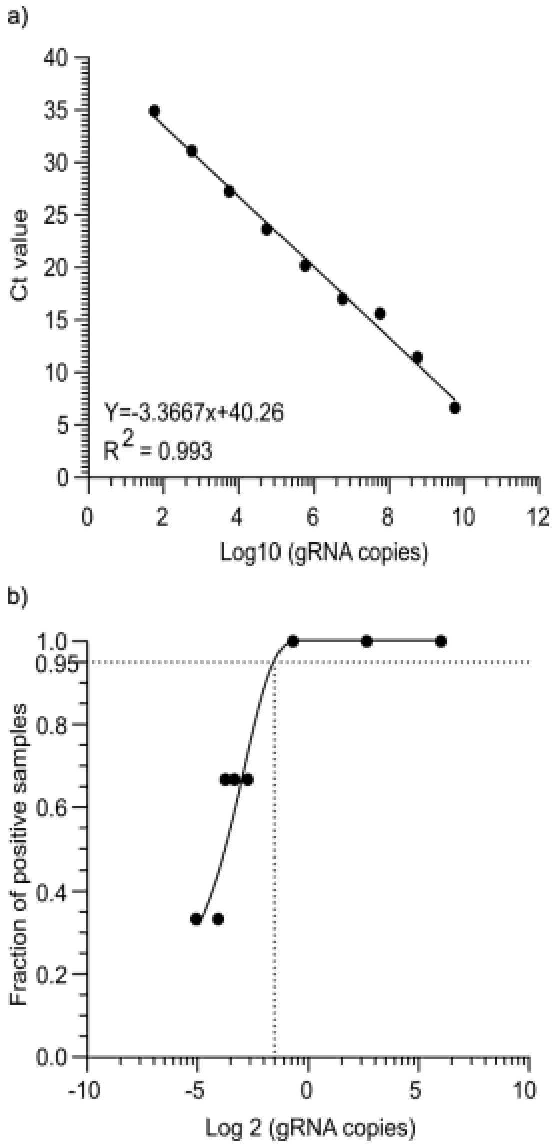

**S8 Fig. (Related to Fig. 1) | Absolute quantification of ZIKV gRNA**.

*In vitro* transcribed RT-qPCR target was serially diluted from 5.75 x $10^9$ to 1 copy before quantification by one-step RT-qPCR. **a** Standard curve for absolute quantification of ZIKV gRNA copies. Points show averaged Ct calculated from three independent replicates. **b** Limit of Detection (LoD) at 95% confidence for ZIKV gRNA is 4 copies ($-1.8^2$). Fractions of positive samples were calculated from three replicates.

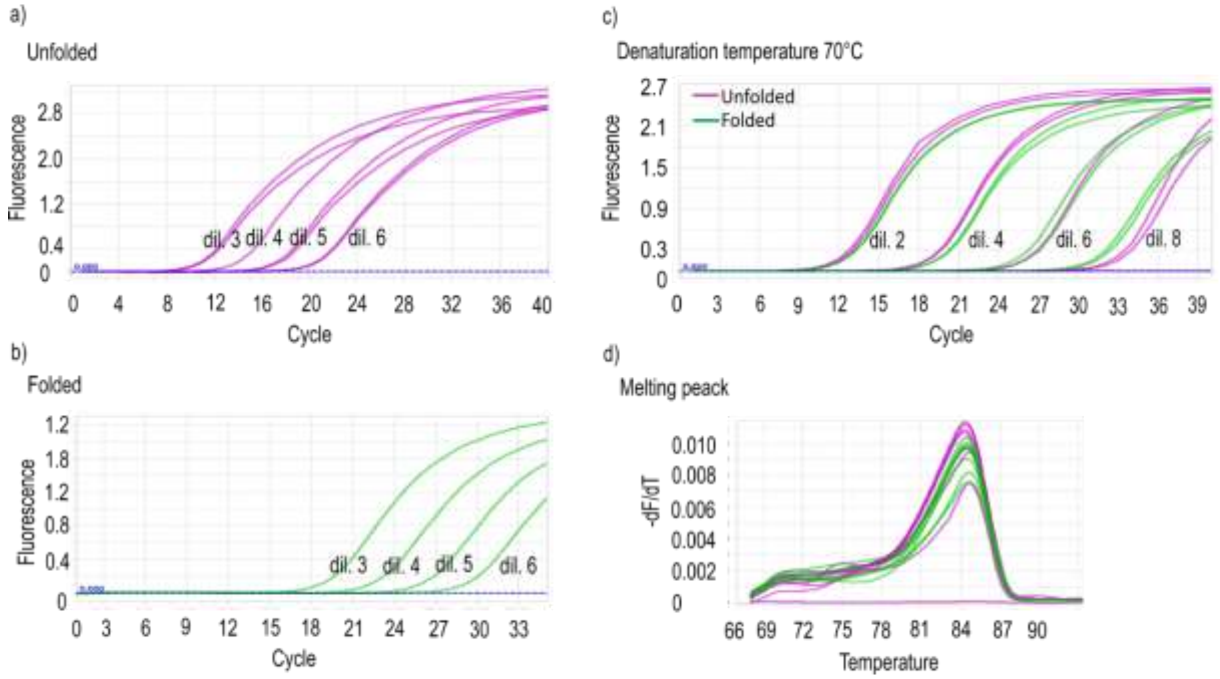

**S9 Fig. (Related to Fig. 1) | Optimization of RT-qPCR quantification for folded ZIKV sfRNA**.

**a-b** Amplification curves for different dilutions (dil. 3, 4, 5, 6) of unfolded (a) and folded (b) *in vitro*-transcribed ZIKV sfRNA in one-step RT-qPCR with RT temperature at 50°C. **c** Amplification curves for different dilutions (dil. 2, 4, 6, 8) of unfolded and folded *in vitro*-transcribed ZIKV sfRNA in two-step RT-qPCR with RT at 70°C. **d** Melting curve analysis for the reaction in c.

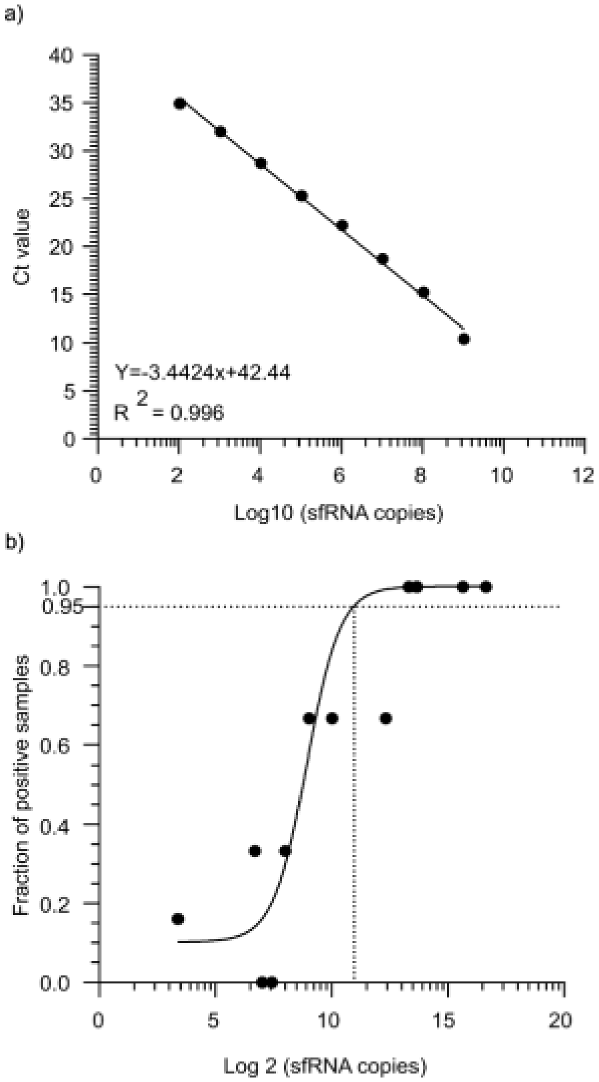

**S10 Fig. (Related to Fig. 1) | Absolute quantification of ZIKV sfRNA**.

*In vitro* transcribed RT-qPCR target was serially diluted from 1.05 x $10^9$ to 10 copies, before quantification by two-step RT-qPCR. **a** Standard curve for absolute quantification of ZIKV sfRNA copies. Points show averaged Ct calculated from three independent replicates. **b** Limit of Detection (LoD) at 95% confidence for ZIKV sfRNA is 152 copies (-$12.3^2$). Fractions of positive samples were calculated from six replicates.

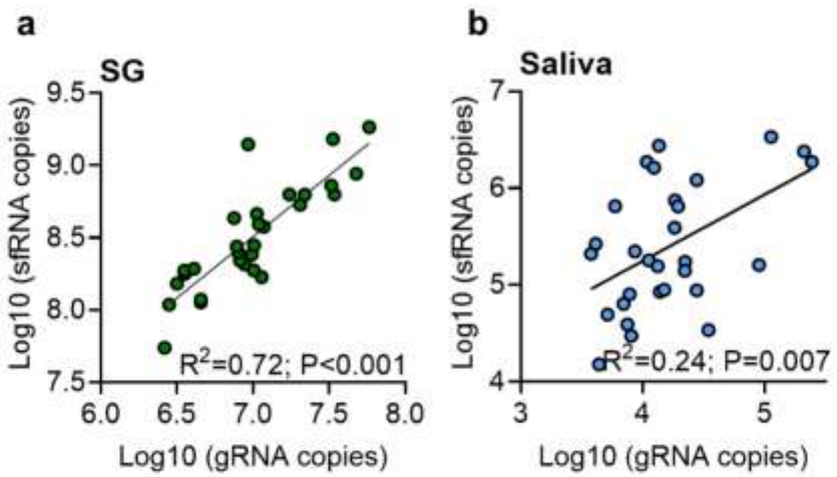

**S11 Fig. (Related to Fig. 1) | a-b Correlations between gRNA and sfRNA copies within salivary glands (SG) and saliva from ZIKV-inoculated mosquitoes**.

Dots indicate SG and saliva repeats. P values were determined by correlation analysis.

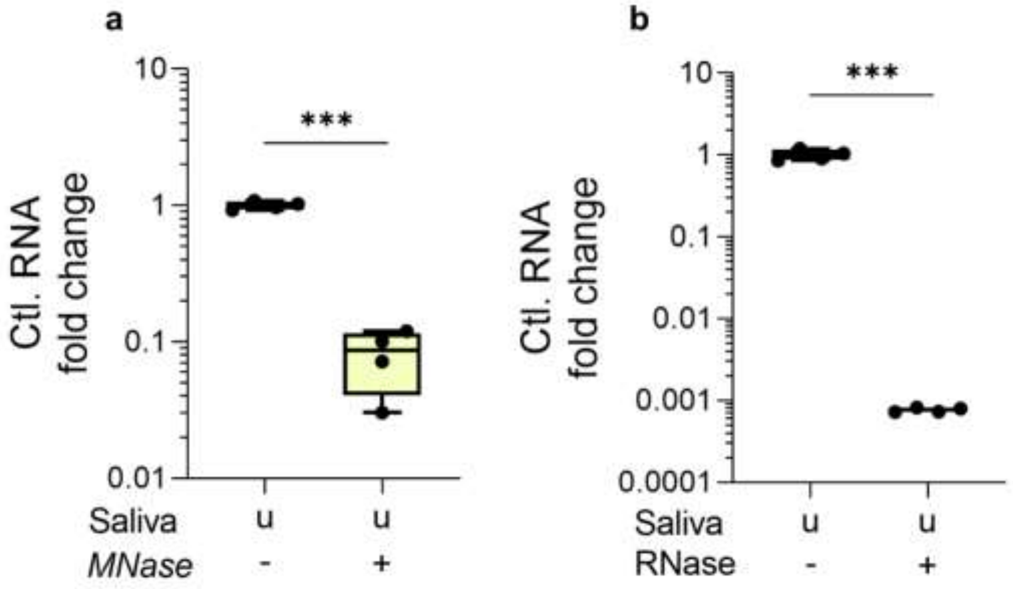

**S12 Fig. (Related to Fig. 2) | a-b MNase and RNase degrade a viral RNA control**.

Boxplots indicate median ± lower and higher quartiles. N, 4. ***, $p < 0.001$ according to T-test.

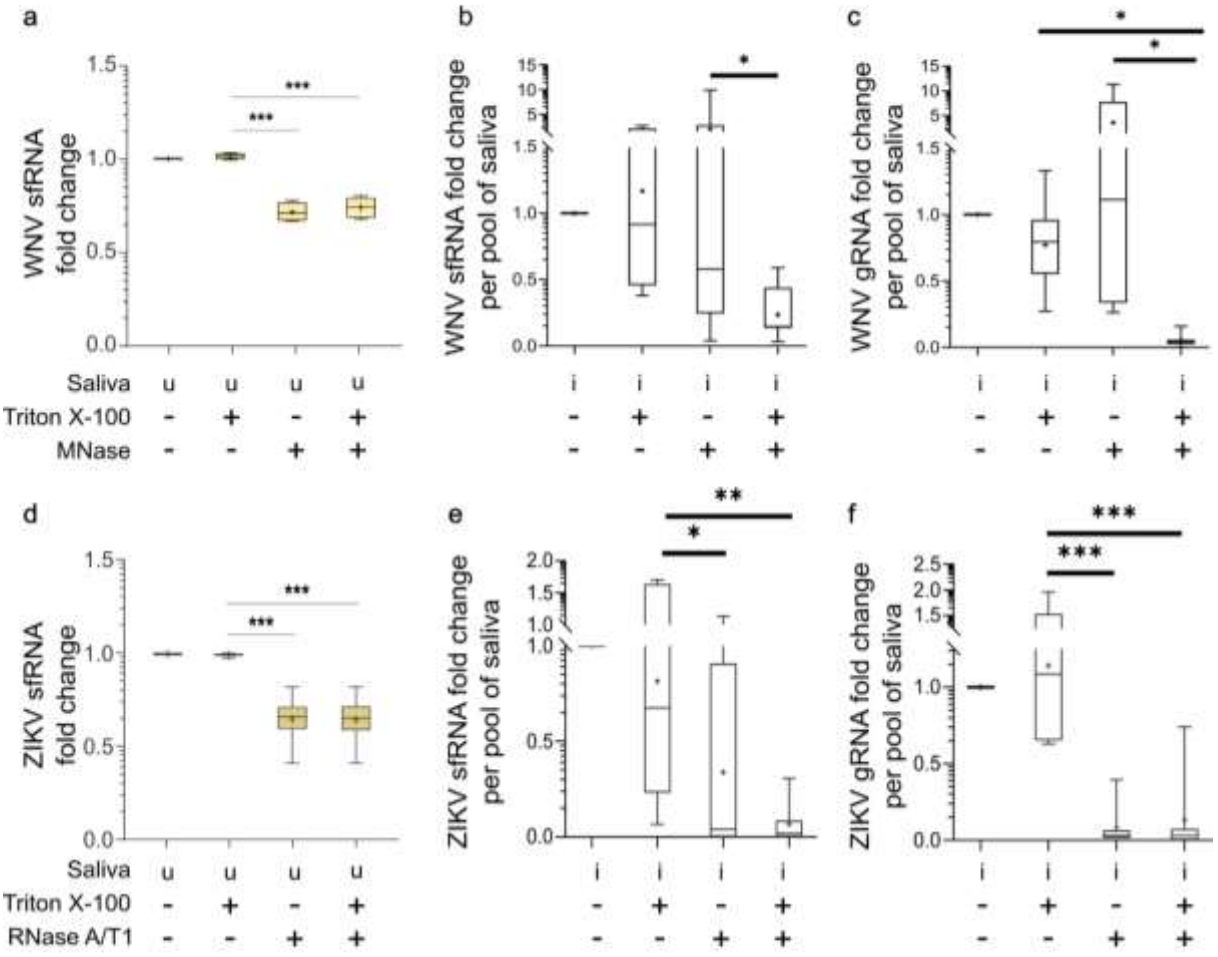

**S13 Fig. (Related to Fig. 2) | Sensitivity to nuclease degradation after detergent treatment for WNV sfRNA in *Culex* saliva and ZIKV sfRNA in *Aedes* saliva**.

The data presented are the same as in fig. 2 except that copies of gRNA and sfRNA were normalized within saliva pools. **a-c** Sensitivity to triton X-100 and Micrococcal nuclease (MNase) treatment for *in vitro*-transcribed folded WNV sfRNA diluted in uninfected *Culex* saliva (a), for salivary WNV sfRNA (b) and salivary WNV gRNA (c) collected from inoculated *Culex* mosquitoes. **d-f** Sensitivity to triton X-100 and RNase A/T1 treatment for *in vitro*-transcribed folded ZIKV sfRNA diluted in *Aedes* uninfected saliva (d), for salivary ZIKV sfRNA (e) and salivary ZIKV gRNA (f) collected from inoculated *Aedes* mosquitoes. Boxplots indicate median ± lower and higher quartiles. N, 8 for WNV and 7 for ZIKV. u, uninfected saliva; i, infected saliva from the corresponding mosquito. *, $p < 0.05$; **, $p < 0.01$; ***, $p < 0.001$ according to T-test.

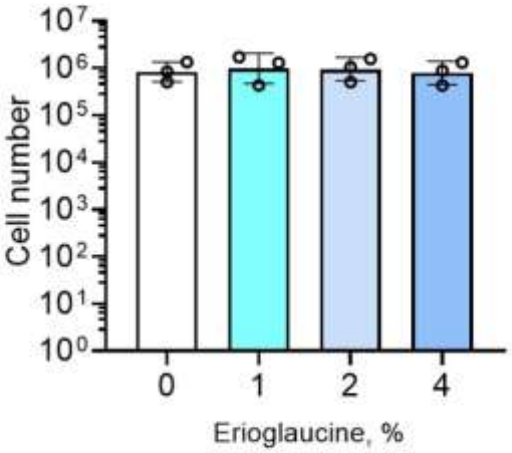

**S14 Fig. (Related to Fig. 3) | Erioglaucine effect on cell survival**.

Number of live cells after 48h of supplementation with 0-4% Erioglaucine. Points indicate repeat. Bars show geometric means ± 95% C.I.

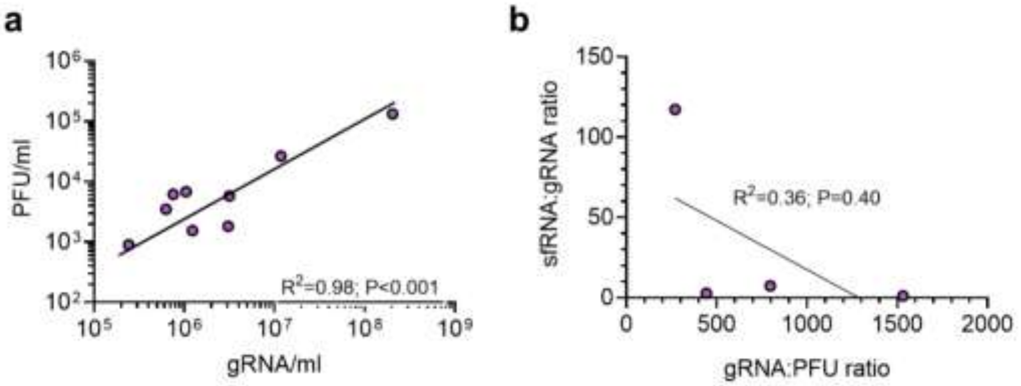

**S15 Fig. (Related to Fig. 3) | Relations between gRNA, PFU and sfRNA in saliva from orally WNV infected mosquitoes**.

**a** Correlation between gRNA and PFU. **b** Correlation between sfRNA concentration (i.e., sfRNA:gRNA ratio) and infectivity (i.e., gRNA:PFU ratio). Linear regression is indicated by the straight line.

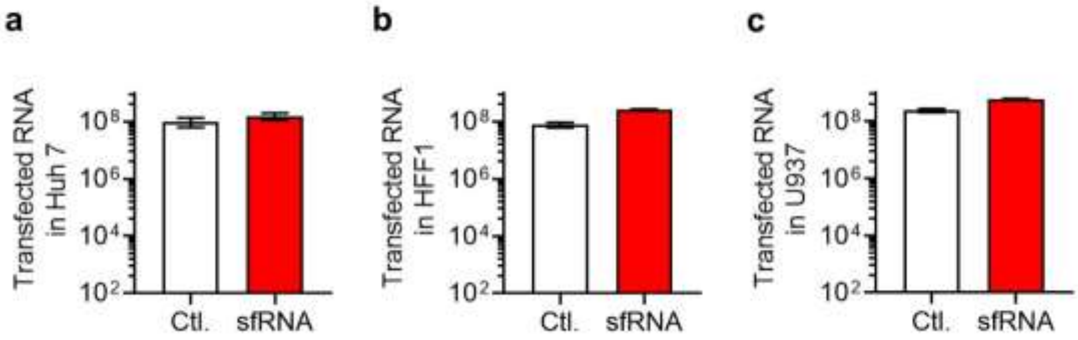

**S16 Fig. (Related to Fig. 3) | Quantification of sfRNA and control RNA transfected in Huh7 (a), HFF1 (b) and U937 (c) cells**.

Cells were transfected with either sfRNA or control RNA (Ctl.) for 2 h and intracellular amount of the respective RNAs were quantified. Bars indicate geometric means ± 95% C.I. from at least three repeats.

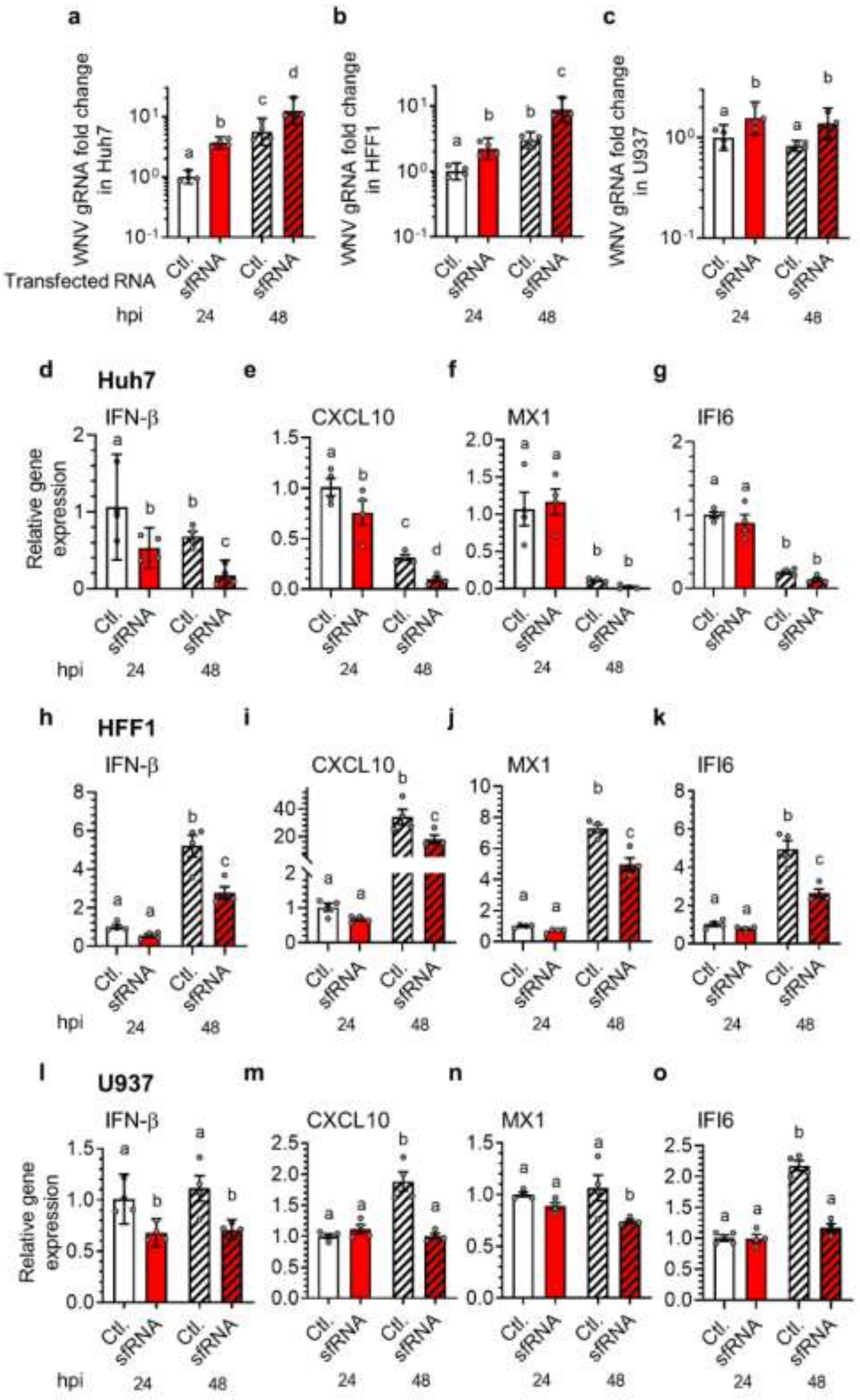

**S17 Fig. (Related to Fig. 3 and 4) | Transfection of sfRNA increases infection and reduces IFN and ISG activation in Huh7, HFF1 and U937 cells**.

**a-c** WNV level after infection at MOI 0.5 in Huh7 (a), HFF1 (b) and U937 (c) cells at 24 and 48 hpi with WNV post sfRNA transfection. **d-g** Expression of IFN-β (d), CXCL10 (e), MX1 (f), and IFI6 (g) in Huh7 cells at 24 and 48 hpi with WNV at MOI 0.5 post sfRNA transfection. **h-k** Expression of IFN-β (h), CXCL10 (i), MX1 (j), and IFI6 (k) in HFF1 cells at 24 and 48 hpi with WNV at MOI 0.5 post sfRNA transfection. **l-o** Expression of IFN-β (l), CXCL10 (m), MX1 (n), and IFI6 (o) in U937 cells at 24 and 48 hpi with WNV at MOI 0.5 post sfRNA transfection. Ctl., RNA control. Repeats are represented by dots. Different letters show significant differences according to post hoc Fisher's LSD test or T-test. a-c Bars show geometric mean ± 95% C.I. n Lines show mean ± s.e.m. d-o Bars show mean ± sem.

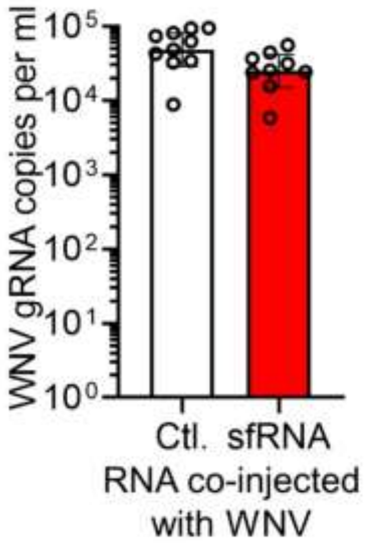

**S18 Fig. (Related to Fig. 3) | Mice injected with either WNV + Ctl. RNA or WNV + sfRNA were successfully infected**.

Blood was collected at 4 days post injection and WNV gRNA copies were quantified per ml of blood. Bars indicate geometric means ± 95% C.I. Repeats are indicated by dots.

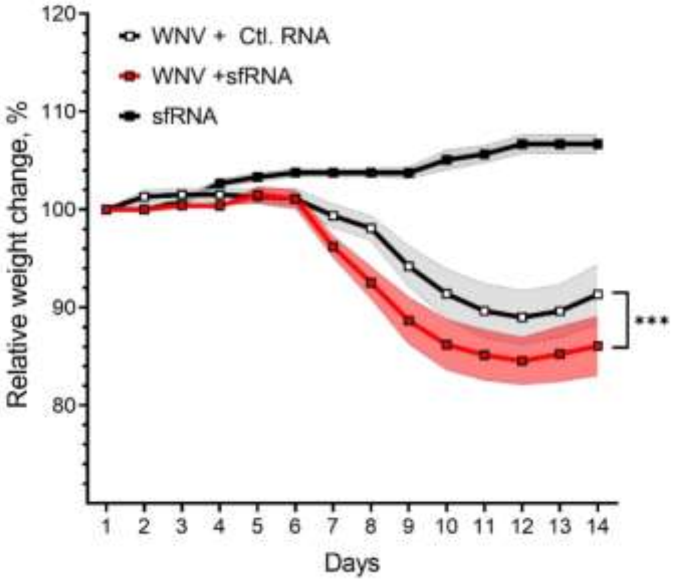

**S19 Fig. (Related to Fig. 3) | Mouse weight change following injection with WNV + Ctl. RNA, WNV + sfRNA or sfRNA alone**.

Lines show mean ± s.e.m. N, 14. ***, $p < 0.001$ according to general linear mixed model followed by Tukey's HSD post hoc test.

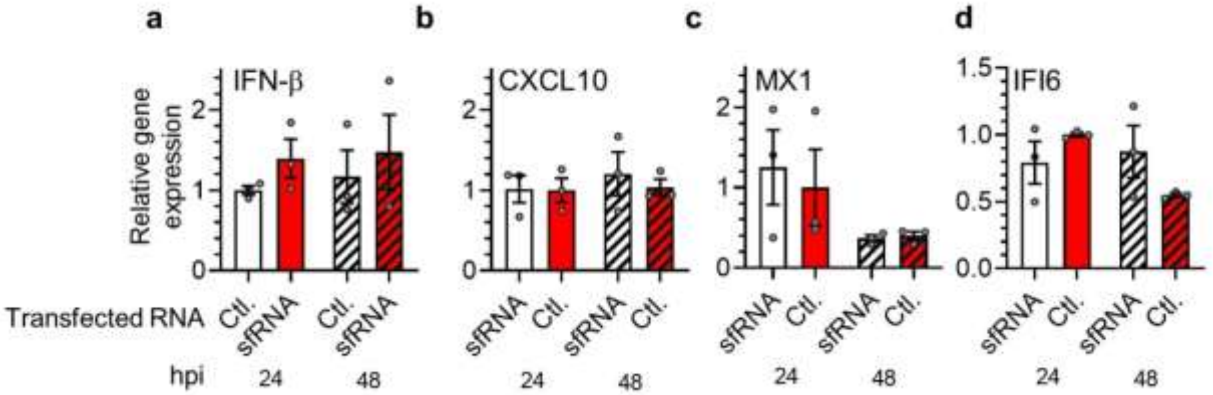

**S20 Fig. (related to Fig. 5) | MRT67037 treatment inhibits the interferon response**.

**a-d** Expression of IFN-β (a), CXCL10 (b), MX1 (c), and IFI6 (d) in Huh7 cells at 24 and 48 hpi with WNV after sfRNA transfection upon MRT67037 treatment. Ctl., RNA control. Bars show mean ± sem. Repeats are indicated by dots.

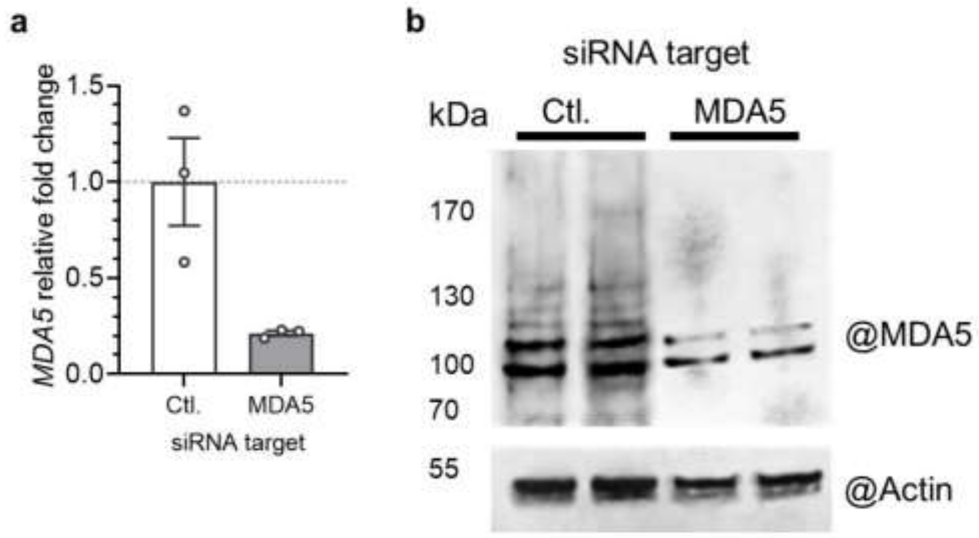

**S21 Fig. (related to Fig. 5) | Silencing of MDA5 in Huh 7.5 cells**.

**a-b** Relative gene expression (a) and protein level (b) of MDA5 at 48 h post siRNA transfection. Actin was used as loading control. MM, molecular markers. Ctl., all stars negative control siRNA were transfected as control.

**S1 Table. (Related to Fig. 3 and 4) | Details of the saliva inoculum used to infect Huh 7 cells**.

Each well was inoculated with saliva containing 1,000 copies of gRNA. The volume of saliva was completed to 50 µl in each well with uninfected saliva.

| Date of saliva pool collection | Cat. of sfRNA concentration | gRNA (copies/µl) | sfRNA (copies/µl) | sfRNA:gRNA ratio | Volume of infected saliva/well (µl) | Volume of uninfected saliva/well (µl) |
|---|---|---|---|---|---|---|
| 07-02-23 | High | 31.76 | 31,926.77 | 1,005.13 | 31.48 | 18.52 |
| 07-02-23 | High | 17.13 | 12,581.69 | 734.48 | 58.38 | 0.00 |
| 09-03-23 | High | 30.42 | 19,475.28 | 640.20 | 33.00 | 17 |
| 23-02-23 | High | 84.73 | 42,477.71 | 501.33 | 11.80 | 38.20 |
| 03-05-23 | Moderate | 34.49 | 8,719.42 | 252.81 | 28.99 | 21.01 |

| | | | | | | |
|---|---|---|---|---|---|---|
| 03-05-23 | Moderate | 35.09 | 7,556.38 | 215.36 | 28.50 | 21.50 |
| 09-03-23 | Moderate | 474.19 | 14,857.19 | 31.33 | 2.11 | 47.89 |
| 09-03-23 | Moderate | 804.25 | 14,920.42 | 18.55 | 1.24 | 48.76 |
| 24-03-23 | Low | 1,124.86 | 5,010.53 | 4.45 | 0.88 | 49.12 |
| 23-02-23 | Low | 22.16 | nd | 0.00 | 45.13 | 4.87 |
| 09-03-23 | Low | 398.08 | nd | 0.00 | 2.51 | 47.49 |
| 24-03-23 | Low | 236.32 | nd | 0.00 | 4.23 | 45.77 |

nd, not detected.

**S2 Table. (Related to Fig. 3 and 4) | Details of the saliva inoculum used to infect human skin explants**.

Each skin explant was inoculated with saliva containing 760 copies of gRNA. The volume of saliva to inject was completed to 30 µl with uninfected saliva.

| Cat. of sfRNA concentration | gRNA (copies/µl) | sfRNA (copies/µl) | sfRNA:gRNA ratio | Volume of infected saliva/well (µl) | Volume of uninfected saliva/well (µl) |
|---|---|---|---|---|---|
| High | 20.20 | 14,189.37 | 702.57 | 37.6 | 0 |
| High | 25.33 | 13,654.49 | 539.11 | 30 | 0 |
| High | 28.27 | 11,797.50 | 417.36 | 26.9 | 3.1 |
| Low | 1239.12 | 9,246.12 | 7.46 | 6.13 [1:10] | 23.87 |
| Low | 39.83 | nd | 0 | 19.1 | 10.9 |
| Low | 67.33 | nd | 0 | 11.3 | 18.7 |

nd, not detected.
